# Exploring protein duplication across Archaea genomes

**DOI:** 10.64898/2026.10.09.758139

**Authors:** Israel Muñoz-Velasco, Valentina Herrera-Cacho, Gerardo Mejia, Angelica Jara-Servin, Leonardo Martinez-Trueba, Luis D. Alcaraz

## Abstract

Gene duplication is a major driver of genomic innovation, but whether paralog accumulation scales uniformly with increasing genome size across the domain Archaea remains unclear. This study presents a domain-wide analysis of 1,829 archaeal genomes and their predicted proteomes to evaluate protein family expansions, their distribution, and lineage-specific patterns of gene duplication. Protein family expansions were classified as recent (90% sequence identity), intermediate (70%), or old (40%) duplications based on sequence-identity thresholds within each family. The findings show that gene duplication content is decoupled from overall proteome size at high sequence-identity levels, indicating that recent paralog retention does not correlate with proteome expansion. Archaeal lineages exhibit contrasting gene duplication profiles. All 1,829 genomes contained paralogs at the 40% sequence identity threshold, indicating the widespread presence of ancient or divergent duplication events. In contrast, paralogs detected at the 90% identity threshold, showed substantial variation across lineages. Their proportion reached up to 16.6% in groups such as Halobacteria, whereas only 74 genomes (4.04%) completely lacked paralogs at this threshold. Highly duplicated families are enriched for membrane transport systems (ABC/MFS), central metabolic enzymes (*e.g.*, acetyl-CoA synthetase), a high proportion of uncharacterized functions, and mobile elements, predominantly transposases. Notably, we identified 314 transposase protein families in Archaea, distributed in 161 unique families in Archaea and 153 families shared with Bacteria, suggesting mechanisms of interdomain gene transfer. Overall, paralog accumulation in Archaea is a lineage-specific phenomenon shaped by targeted functional duplications and mobile element activity, rather than a simple consequence of proteome growth.

**Significance:** It has been uncertain whether high levels of gene duplications across archaeal genomes represent proportional genome growth or specific expansions. Our analysis of reference archaeal proteomes reveals that gene duplication is decoupled from genome length, as high-identity duplications do not scale linearly with proteome size. Instead, archaeal lineages exhibit different duplication profiles, including small-genome organisms with up to 18.3% of their proteome to paralogous groups, as well as specific clades displaying duplications of transport systems, central metabolic enzymes, and transposases. Together, these findings show that gene duplication content in Archaea is shaped by targeted functional expansions and mobile genetic element activity rather than overall proteome size.

## 1. Introduction

The evolutionary success of the Domain Archaea, characterized by their ability to thrive in the most extreme niches on Earth, including hypersaline basins, hydrothermal vents, and other environments, is fundamentally linked to their genomic plasticity. A key element of this adaptability is genome expansion, driven by the duplication of genes, metabolic modules, and sometimes entire genomic segments (Pecoraro et al. 2011; Soppa 2017). Gene duplication is a key driver of innovation, providing the genetic material for functional diversification, the development of new traits, and increased resilience to environmental changes (Hughes 2005; Copley 2020; Sheridan et al. 2020).

While classical evolutionary models, such as the “Patchwork Model,” propose that ancient metabolic enzymes originated through the recruitment of promiscuous activities followed by gene duplication and subsequent substrate specialization (Jensen 1976; Caetano-Anollés et al. 2009; Fani and Fondi 2009), contemporary archaeal genomics reveals that duplication is not merely an ancestral vestige of early life; rather, it is a dynamic, ongoing response to environmental change (Tria and Martin 2021). Despite general scaling laws predicting a correlation between total proteome size and gene duplication levels, many archaeal lineages exhibit highly duplicated genomic architectures in which internal duplication levels are decoupled from overall genome size (Martínez-Núñez et al. 2013; Álvarez-Lugo and Becerra 2023). This implies that specific lineages prioritize expanding functional modules, particularly those related to transcriptional regulation and stress-response pathways, as an adaptation to ecological constraints (Edgell et al. 1998; Álvarez-Lugo and Becerra 2021).

Furthermore, the complex interplay between the mobilome and core genomic architecture suggests that transposable and extrachromosomal elements actively catalyze evolution (Archibald and Roger 2002; Manzano-Morales and Gabaldon 2025). These elements maintain high-identity paralogs that facilitate rapid adaptive rearrangements, which are very important for lineages inhabiting fluctuating environments (Edgell et al. 1997; Hooper 2003). Previous studies have attempted to correlate ancient, highly divergent paralogs with recent, conserved duplications, thereby elucidating the evolutionary forces that shape new traits (Vosseberg et al. 2020; Tria et al. 2021).

In this study, we analyze gene duplication patterns across 1,829 selected complete archaeal genomes by integrating phylogenetic information with annotation and functional ontology profiling, characterizing how duplication profiles diverge across taxonomic classes, from streamlined lineages to those with extensively duplicated genomes, and demonstrating that functional traits structure these profiles. Furthermore, we characterize transposable elements as critical drivers of genomic architectural plasticity; by performing a large-scale cross-domain screening of transposases, we reveal numerous archaea-specific and shared mobile genetic elements in their genomes. Through this study, we highlight the interplay between ancient and recent duplications, providing a comprehensive view of whether a highly duplicated genomic architecture serves as an architectural trait or a specialized survival strategy across different environments.

## 2. Results

### 2.1 Global patterns of gene duplication across Archaea

To obtain a general overview of proteome sizes, we analyzed the protein count across 1,829 archaeal proteomes. The distribution revealed high heterogeneity, appearing to follow a bimodal pattern with two distinct peaks, one centered at approximately 2,500 proteins, exceeding 150 genomes, and a second peak near 3,500 proteins, encompassing over 130 genomes. Most genomes ranged between 2,000 and 4,500 proteins, though significant tails extend toward highly reduced and expanded states (Figure 1a). This variation in proteome size provides the necessary background to evaluate whether gene duplication scales uniformly across the domain or is driven by lineage-specific evolutionary trajectories.

**Fig. 1.**
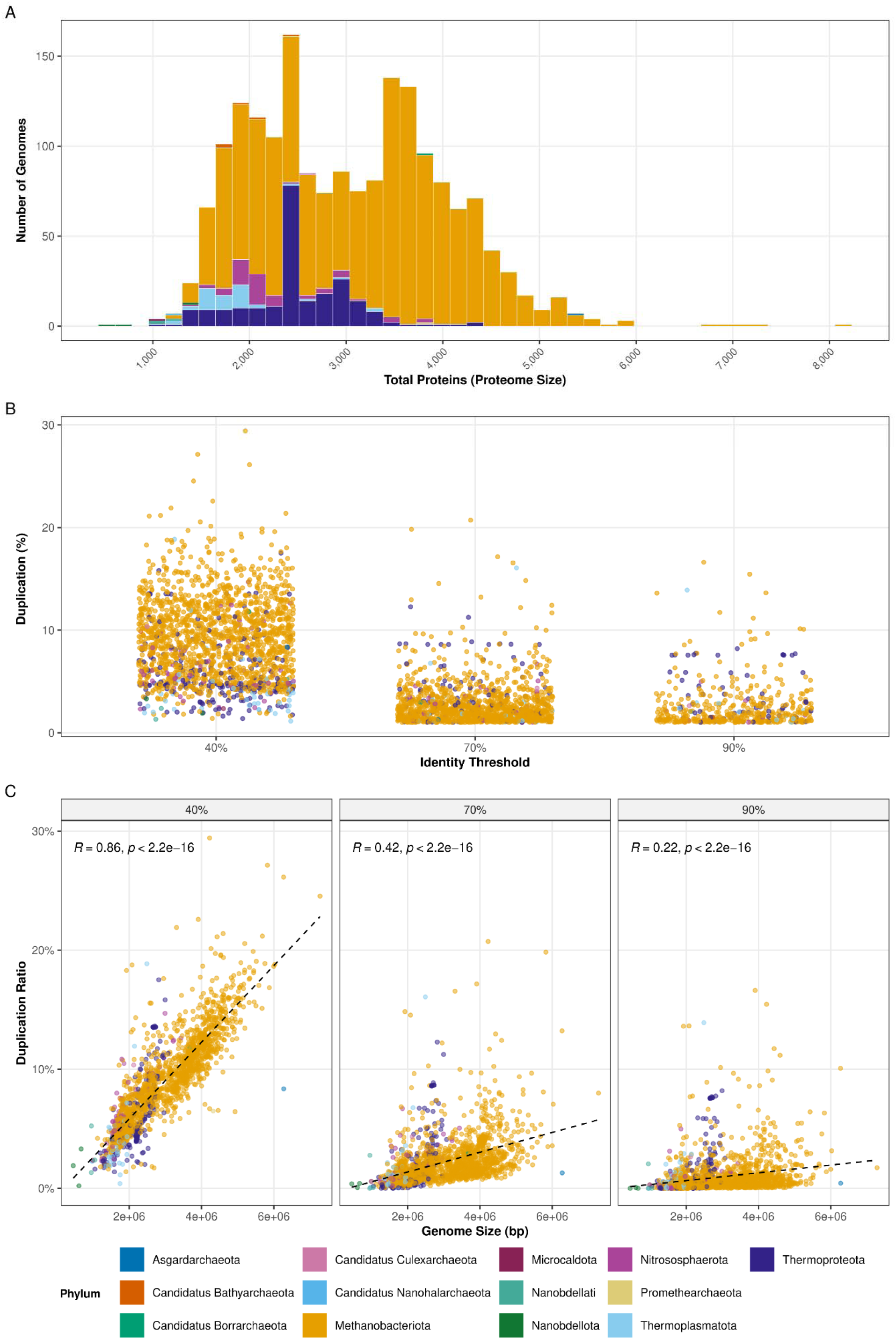
Global distribution of proteome sizes and gene duplication levels across 1,829 archaeal genomes. (a) Histogram of total protein-coding genes (proteome size) per genome. (b) Distribution of the absolute count of duplicated proteins identified at three sequence identity thresholds (40%, 70%, and 90%). Dots represent individual genomes with more than one duplicated protein. (c) Paralogous gene content expressed as a percentage of the total proteome across the same identity thresholds. Each facet illustrates the relative proportion of the proteome composed of paralogous groups. Points represent individual genomes with duplication levels above 1%. Each color represents a different Phylum within the Archaea Domain. **Alt text.** Three plots showing archaeal proteome distribution. Panel a is a histogram of protein-coding gene counts per genome. Panel b is a scattered plot of duplicated protein counts across identity thresholds. Panel c shows paralog percentage across identity thresholds. Data points are colored by phylum.

Our results showed that gene duplication is a universal feature of the domain Archaea. At the most permissive evolutionary threshold (40% identity), which captures both ancient duplications and diversified gene families, every genome in the dataset possessed at least one pair of paralogous proteins, resulting in zero genomes without duplicated genes. At 40% identity, the paralogous repertoire was substantial, with a median of 8.71% of the proteome assigned to duplicated families (Interquartile Range: 5.74% – 11.50%) (Table 1). At this threshold, the paralogous distribution is characterized by extensive genome-wide dispersion, with a significant proportion exhibiting elevated duplication densities, where paralogous content frequently exceeds 5% and reaches a maximum of 29.4% (Figure 1b). This widespread presence suggests that highly divergent duplications are not merely random events but a core structural component of the archaeal genomic framework.

**Table 1.** Summary statistics of duplicated proteome fraction (%) across 1,829 archaeal genomes at three identity thresholds.

| Threshold | Median Duplicated Proteome (%) | Q1 (25%) | Q3 (75%) | Max | Genomes Without Duplications* |
| --- | --- | --- | --- | --- | --- |
| 40% | 8.71 | 5.74 | 11.50 | 29.40 | 0 |
| 70% | 1.62 | 0.93 | 2.73 | 20.70 | 5 |
| 90% | 0.47 | 0.18 | 1.06 | 16.60 | 74 |

\*Genomes without duplicated proteins detected at that specific identity threshold.

We observed a rapid decay of the duplication signal as sequence-identity stringency increased. When transitioning from the 40% threshold to the 70% threshold, we observed a dramatic drop in median paralogous content, from 8.71% to 1.62% (Table 1). This decay continues toward the 90% threshold, where the median levels off at 0.47%. This pattern also appears in the absolute counts of duplicated proteins (Figure 1b), where most genomes shifted from hundreds of divergent paralogs at 40% identity to fewer than 10 proteins retained in highly identical paralogous families at 90% identity (see Figure 1b and Table 2).

**Table 2.**
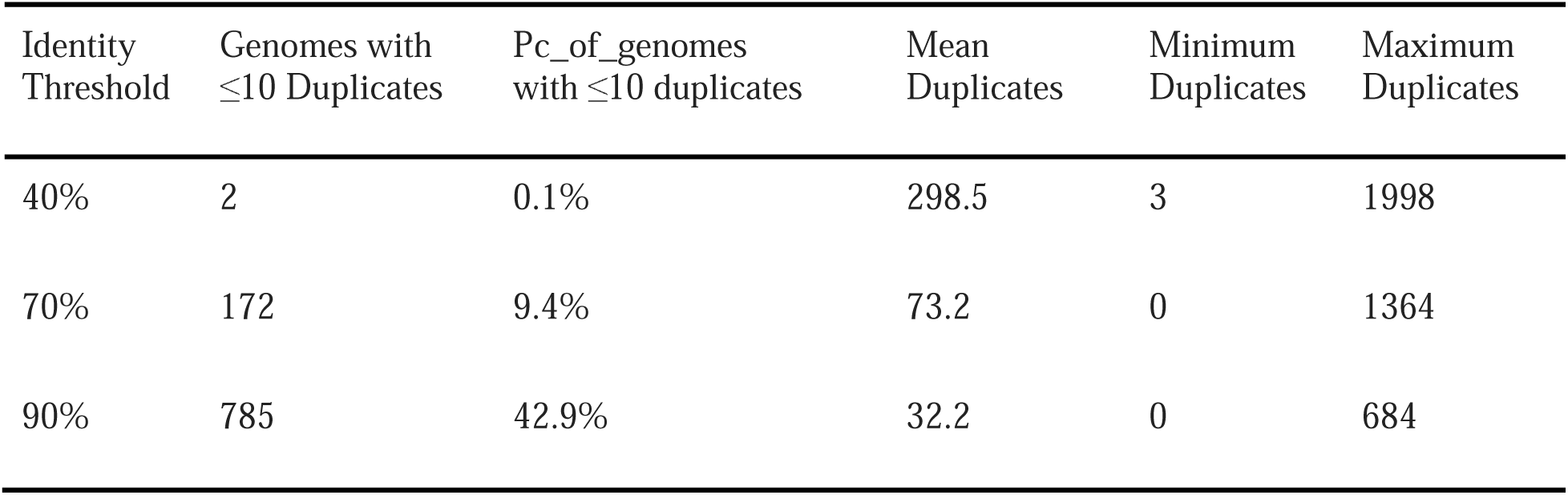
Duplication statistics across identity thresholds for 1,829 Archaea genomes. Values represent protein family duplicates.

| Identity Threshold | Genomes with $\leq 10$ Duplicates | Pc_of_genomes with $\leq 10$ duplicates | Mean Duplicates | Minimum Duplicates | Maximum Duplicates |
| --- | --- | --- | --- | --- | --- |
| 40% | 2 | 0.1% | 298.5 | 3 | 1998 |
| 70% | 172 | 9.4% | 73.2 | 0 | 1364 |
| 90% | 785 | 42.9% | 32.2 | 0 | 684 |

Although the overall proportion of duplicated genes diminished and the distribution stabilized at higher identity values, certain lineages remain notable exceptions to this conserved archaeal architecture. While 74 genomes (4.04%) showed no recent duplications at the 90% threshold (Table 1), we identified 72 paralog-enriched genomes. These outliers, termed “Double Outliers,” ranked in the top 5th percentile for both the absolute number of paralogs and the total percentage of duplicated proteins (Supplementary Table S1), with values reaching 29.4% at the 40% identity threshold (Figure 1b).

Our analysis revealed that extensive gene duplication is not merely a byproduct of genome expansion, because while the majority of duplicated-enriched genomes are indeed large, we identified a specific group of three lineages designated as “Small-Genome High-Duplication” (SGHD) outliers that maintain an average proteome size of approximately 2,495 proteins but exhibited disproportionately high paralogous content, ranging from 18.3% to 18.9% (Supplementary Table S1). These SGHD genomes include *Ferroplasma sp*. Type II (GCF_000496155), *Thermococcus bergensis* T7324 (GCF_020386975), and *Methanotorris formicicus* Mc-S-70 (GCF_047915675), suggesting that in certain archaeal groups, gene duplication is a targeted adaptive strategy that occupies a substantial share of the genomic space, rather than a non-specific consequence of overall genome growth.

At the 40% identity threshold, we observed a strong positive correlation (R = 0.86, R^2^ = 0.74, P < 2.2 x 10^-16^), indicating that proteome expansion explains nearly 74% of the variance in total paralog content (Figure 1c). At the 70% threshold, the correlation coefficient drops to R = 0.42, and by the 90% threshold, the relationship becomes weak (R = 0.22). At this stringent identity threshold, the data form a highly dispersed cloud in which the size dependency effectively disappears. We observed numerous expanded genomes exceeding 4,500 kbp that maintain a low duplication ratio, often below 2.5% at 70% and 90% identity thresholds, indicating that a large genomic template does not inherently lead to retaining identical gene copies. Conversely, we identified smaller lineages (starting near 2,000 kbp) that exhibit high-identity duplication ratios of up to ∼14% at the 90% threshold (Figure 1c).

### 2.2 Identification of outlier genomes and taxonomic structure of duplication patterns

To identify lineages that defy domain-wide trends, we implemented a residual-based detection strategy. By contrasting observed duplication counts with expected values from our linear models across archaeal genomes, we treated deviations from the norm as meaningful biological signals (Supplementary Figure S1). This multi-scale approach allowed us to differentiate genomes that merely follow the scaling law from those that have undergone atypical, lineage-specific expansions. At the 40% identity threshold, we identified 75 genomes (Supplementary Table S1) as significant outliers, in which the duplication levels exceed the predicted threshold of the expected distribution (Z-score > 2), effectively marking these taxa as hotspots of genomic duplications.

The systematic analysis of archaeal genomic architecture reveals that gene duplication in this domain is not homogeneous but is instead tightly linked to phylogenetic heritage and lineage-specific selective pressures. By examining the distribution of duplication density across the 28 identified taxonomic classes, we observe that ancestral paralogous content at the 40% identity threshold fluctuates drastically, ranging from a minimal 1.06% in the most streamlined groups to a substantial 11.8% of the total proteome in the most expanded lineages (see Supplementary Table S2). This taxonomic segmentation suggests that paralog accumulation is not a consequence of proteome growth but an evolutionary strategy differentially adopted across the domain.

The visual inspection of the relationship between duplication density and proteome size, as shown in Figure 2, confirms genomic hotspots associated with specific Phyla. The Class Halobacteria, within the Phylum Methanobacteriota, stands out as the most prominent example of this trend, accounting for a significant portion of all duplication-enriched outliers. Using standardized residuals (Z-scores) as a measure of relative duplication density, Figure 2 shows how different lineages manage their paralogous repertoires across identity thresholds at 40% (Figure 2a) and 90% (Figure 2b), with more representative genomes exhibiting higher Z-scores in Methanobacteriota and Thermoproteota. This comparative approach functions as an evolutionary continuum of gene duplication. We discerned distinct expansion trajectories; for example, some lineages, such as several Halobacteria, exhibit high paralogous content across all identity thresholds (Z > 4), while others show a sharp decay at the 90% threshold (*e.g.*, some members of Nanobdellati and Microcaldota) (Figure 2b).

**Fig. 2.**
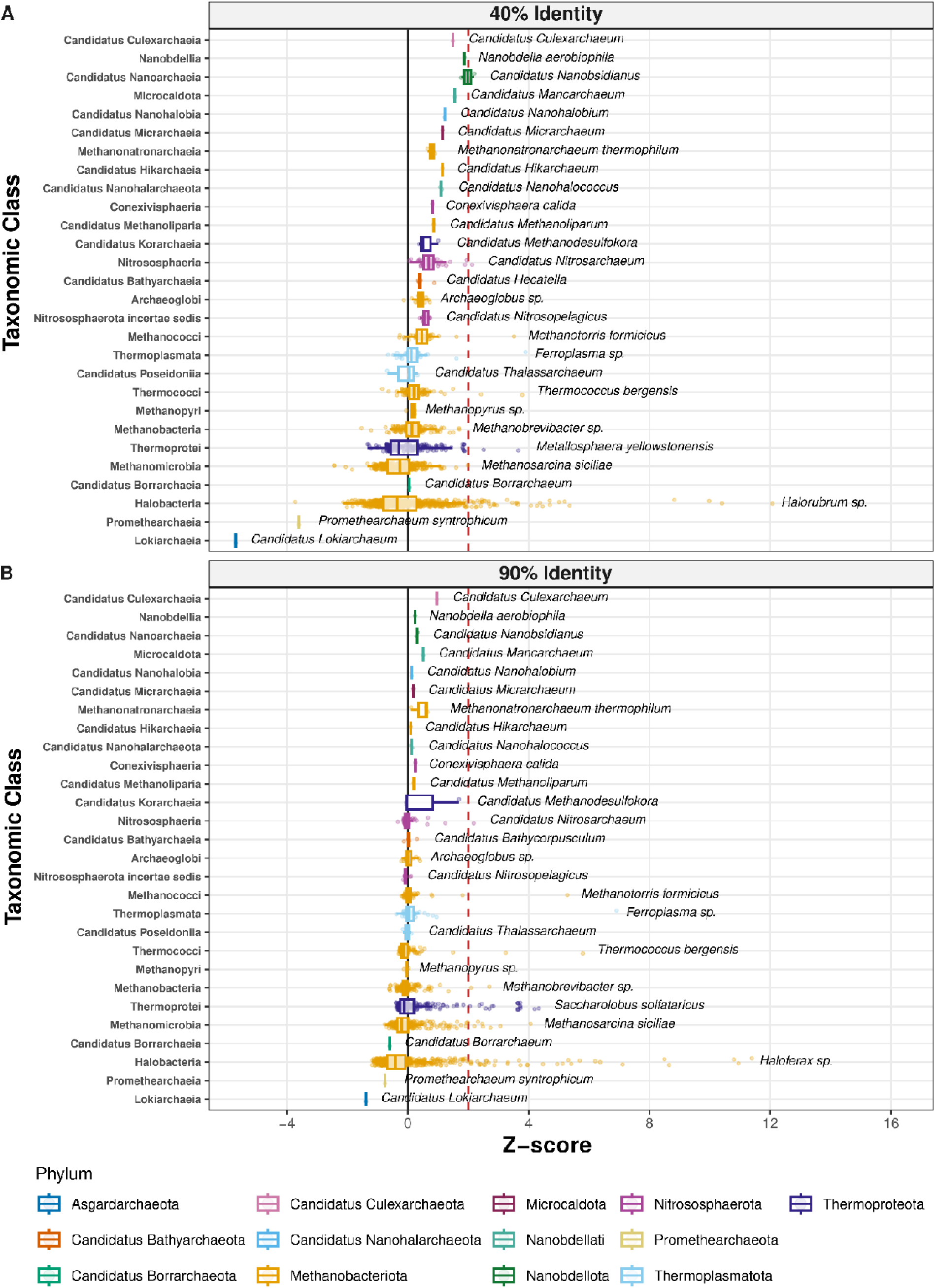
Taxonomic distribution of gene duplication across three evolutionary scales. The boxplots represent the distribution of standardized residuals (Z-scores) derived from the linear scaling relationship between proteome size and gene duplication. Data are organized by taxonomic class (y-axis) and faceted by sequence identity thresholds. (a) 40% identity, representing ancestral/divergent gene family expansions; and (b) 90% identity, capturing recent, highly conserved duplications. The solid vertical line at Z = 0 indicates the expected duplication level based on proteome size, while the red dashed line at Z = 2 denotes the threshold for statistical outliers (atypical expansions). Colored points represent individual genomes, and colors correspond to the respective Phylum for each class. **Alt text.** Boxplots arranged vertically by archaeal taxonomic class showing standardized Z-scores of gene duplication density. Panel a displays values at forty percent sequence identity. Panel b shows values at 90% identity. Points are colored by phylum.

To understand the evolutionary structure of these duplicates, we evaluated their sequence conservation using the Persistence Index (PI), defined as the proportion of recent duplicates ( 90% identity) relative to the total set of detectable duplications ( 40% identity) as follows 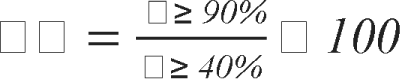. This parameter showed two divergent evolutionary trajectories across Archaea. Certain lineages displayed an active expansion profile of duplicates, with a significant fraction of paralogs remaining nearly identical. These cases of high duplication persistence are primarily concentrated in specific classes, including Microcaldota (PI = 39.12%) and Methanonatronarchaeia (PI = 31.22%), as shown in Supplementary Table S2. In these groups, the duplication machinery appears fully operational, potentially driven by recent environmental adaptations or by mechanisms such as gene conversion that preserve sequence homogeneity.

Conversely, other lineages showed an absence of recent duplications, exhibiting extremely low persistence. As shown in Supplementary Table S2, classes such as Nanobdellia, C. *Nanohalobia,* and *C. Nanohaloarchaeota* show a Persistence Index of 0%, indicating that, although they harbor ancient duplications detectable at 40% identity, none of these events exceed the 90% threshold.

Notably, despite having the highest number of ancestral outliers, the *Halobacteria* showed moderate persistence (9.41%), reflecting a balanced evolutionary model. This dynamic, summarized alongside all other clades in Supplementary Table S2, ensures an uninterrupted supply of raw genetic material for adaptation in hypersaline environments. This suggests that by maintaining a steady influx of new duplications, these archaea can explore new metabolic functions without losing the stability of their fundamental genomic architecture.

### 2.3 Functional ontology characterization of expanded gene families

The functional ontology characterization of duplications identified in duplicated-enriched outliers annotated with eggNOG-mapper (see Material and Methods section) revealed targeted enrichment of biological processes. Our analysis illustrates the distribution of duplicated genes, across sequence identity thresholds of 40% and 90%, passing through the central COG supercategories (mapped from eggNOG descriptions), including mobile elements segregated via keyword regular expressions (see Functional annotation section of Material and Methods), and showing a significant fraction of “functional dark matter” within the Poorly Characterized categories (Figure 3a), suggesting the existence of unique archaeal adaptation mechanisms yet to be discovered. These duplications represent high-priority targets for future experimental characterization to fully decode the ecological success of highly duplicated archaeal lineages.

**Fig. 3.**
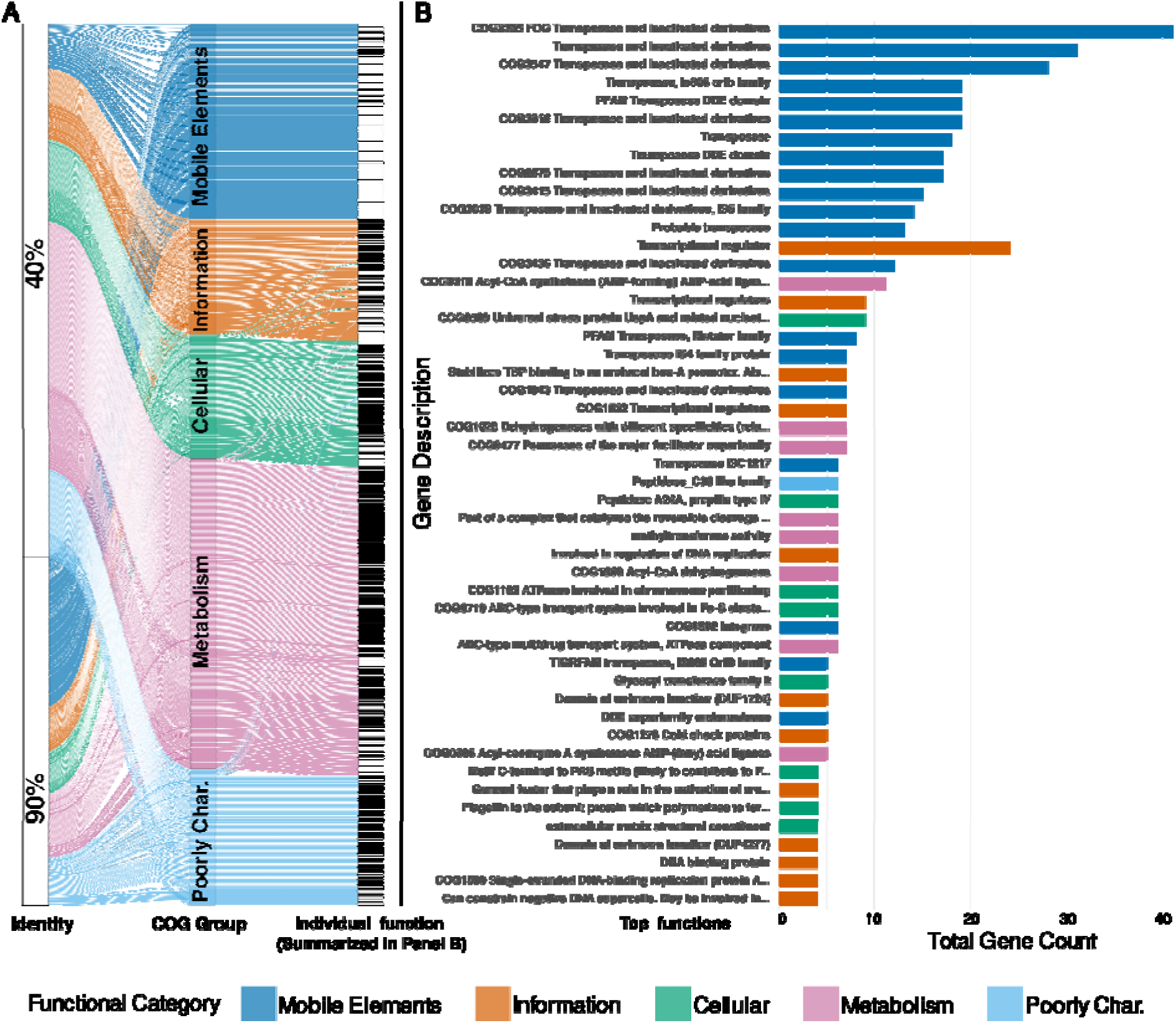
Functional flow of archaeal gene duplications. (a). The alluvial plot illustrates the distribution and transition of expanded gene families from their identity thresholds (40% and 90%) through COG supercategories to their specific functional annotations (individual functional flows are resolved and quantified in panel b. Mobile genetic elements (*e.g.*, transposases and integrases) were identified by regular expressions and separated into a dedicated category alongside standard COG classes. (b). Rank-abundance of the top 50 most frequent specifi functional descriptions derived from the alluvial functional stream. **Alt text.** Two-panel plot. Panel a is an alluvial diagram illustrating gene family transitions from sequence identity thresholds to COG functional categories. Panel b is a horizontal bar chart listing top functional descriptions for duplicated elements.

The top specific Functional Descriptions derived from the eggNOG-mapper analysis showed an important enrichment of transposase-annotated functions (Figure 3b). This analysis showed that Information Storage and Processing (specifically Transcriptional Regulation), along with Mobile Elements and Metabolism, are the primary drivers of gene expansion. Furthermore, Supplementary Figure S2 shows that the functional analysis of duplicated genes varies across the different sequence identity thresholds (40%, 70%, and 90%). High Z-scores are observed at recent identity thresholds for category L (Replication, recombination, and repair), indicating that machinery associated with genomic plasticity and mobile elements is a continuously expanding component of the archaeal genome. In contrast, essential information-processing categories, such as J (Translation) and K (Transcription), exhibit heavy shading at the 70% threshold but fade to light or white tones at 90%, suggesting that recent expansions of core informational machinery are strictly constrained (Supplementary Figure S2).

Metabolic pathways, specifically those involved in energy production, amino acid transport, and inorganic ion transport, showed a clear distribution pattern, with the highest number of duplications at the 40% threshold, reflecting an oldest expansion that gradually tapers off at higher identity levels; nevertheless, distinct signals remain visible at the 70% and 90% identity thresholds. Finally, category S (Function unknown) showed that a substantial portion of the actively expanding genetic pool in Archaea remains uncharacterized, with its highest frequency observed at the 90% identity threshold (Supplementary Figure S2).

Transcriptional regulators, the universal stress protein UspA, and various dehydrogenases, such as COG1028 and Acyl-CoA dehydrogenases, represented the primary categories of ancestral expansions (40%) identified across the dataset (Supplementary Table S3). These expansions were characterized by a high frequency of duplications within the Major Facilitator Superfamily (MFS) and ABC-type transport systems, which dominated the duplication profile of these lineages. The 70% identity threshold serves as an evolutionary bridge through which more specific cellular maintenance functions emerge. At this level, components such as Cation transport ATPases and general transcription factors that stabilize TBP binding to the archaeal box-A promoter were the most duplicated (Supplementary Table S3).

Recent duplications, captured at the 90% identity threshold (Recent), showed a distinct pattern oriented toward genome protection and immediate environmental response. There was a marked enrichment of cold shock proteins, DNA-binding replication proteins (RPA), enzymes that maintain genomic integrity at high temperatures, and a high frequency of Peptidase A24A (prepilin type IV) and ABC-type transport systems involved in Fe-S cluster assembly (see Supplementary Table S3). At the same time, Defense Mechanisms, Signal Transduction Mechanisms, Cell Motility/Post-translational Modification of Proteins and Chaperones, Replication, Recombination and Repair Mechanisms emerged with high values (Supplementary Figure S2) at this threshold.

By mapping KEGG orthology (KO) across distinct identity thresholds, we observed that the 40% identity cut-off is dominated by ABC-2-type transport systems, transcriptional initiation factors such as TFIIB (K03124), and key enzymes for central carbon flux, including acetyl-CoA synthetase (K01895) and acyl-CoA dehydrogenase (K00249). As sequence identity increased to 70%, there was an enrichment of high-affinity metal transporters, such as P-type Cu^2+^ ATPases (CopB) and Zn^2+^/Cd^2+^ exporters (ZntA), and protein-quality-control chaperones, such as VCP/CDC48 (K13525). The 90% identity category was biased toward cold-shock protein A (CspA), a specialized suite of methylamine-corrinoid methyltransferases (K16176–K16178), DNA-binding protein 7 (Ssh7), some resolvases (K07450), and specialized carbon fixation subunits such as cdhE (K00197) (Supplementary Table S4). A core set of functions, most notably acetyl-CoA synthetase (K01895) and the transcription factor TFIIB (K03124), exhibited persistent duplication across all identity thresholds.

### 2.4 Contribution of transposases to paralogous expansion

Given the notable prevalence of Mobile Genetic Elements (MGEs) and transposase-annotated functions observed in our functional profiling (Figure 3b), we performed a systematic cross-domain screening of transposases (Tnps) to evaluate their specific contribution to paralogous diversification across archaeal genomes. We constructed a specialized reference database by clustering identified Tnps (see Materials and Methods, Transposase database construction and distribution analysis section), yielding a curated repertoire of 2,034 representative Tnp families. We first analyzed the distribution of Tnp families across the RefSeq Archaeal genomes and a representative subset of bacterial genomes from the iTOL dataset. Later, by analyzing the full RefSeq bacterial database alongside the 1,828 archaeal genomes, we observed that the domain-specific distribution of these elements underscores a clear distinction in the mobilome between the two domains.

While 1,720 families were exclusive to Bacteria, reflecting the vast diversity of Insertion Sequences (IS) documented in that domain, our analysis identified 161 Tnp families strictly restricted to Archaea and 153 families shared between the two domains (Supplementary Dataset 1 and Supplementary Figure S3). These 314 archaeal-associated families (161 exclusively in Archaea and 153 shared with Bacteria) suggest an independent evolutionary trajectory for MGEs within the archaeal domain. These elements were not uniformly distributed but exhibited distinct patterns of expansion that correlated with specific lineages and lifestyle strategies; this is particularly evident in lineages where extreme environmental pressures, such as high salinity or extreme thermophilicity, may favor the persistence of specialized transposition machinery that was not observed in bacterial counterparts (Supplementary Figure S4).

When we analyzed the number of Tnps duplications across genomes, we observed a non-random distribution. The Principal Coordinate Analysis (PCoA) explained 2.42% of the variance on its first axis and 2.32% on its second axis. Archaea and bacterial genomes showed distinct sets of transposases (PERMANOVA, 999 permutations, R2 = 0.00894, p < 0.01). Archaea showed greater dispersion, with separation along both axes, whereas most bacterial genomes were positioned near the center of the ordination, with separation mainly along the second axis (Figure 4a). Furthermore, transposase family content also differed taxonomically at the phylum level (PERMANOVA, 999 permutations, R2 = 0.0794, p < 0.01), with the most evident separation between Methanobacteriota and Thermoproteota; in particular, Methanobacteriota genomes were distributed into two clusters, oriented along each axis. The remaining phyla were distributed along the second axis: members of Thermoproteota showed little overlap with Methanobacteriota genomes. In contrast, representatives of Thermoplasmatota and Nitrosphaerota showed similarity with members of Methanobacteriota and Thermoproteota (Figure 4b). In Bacteria, the different phyla showed little dispersion among themselves, indicating that the set of transposases shared with Archaea is similar across multiple bacterial lineages (Supplementary Figure S5).

**Fig. 4.**
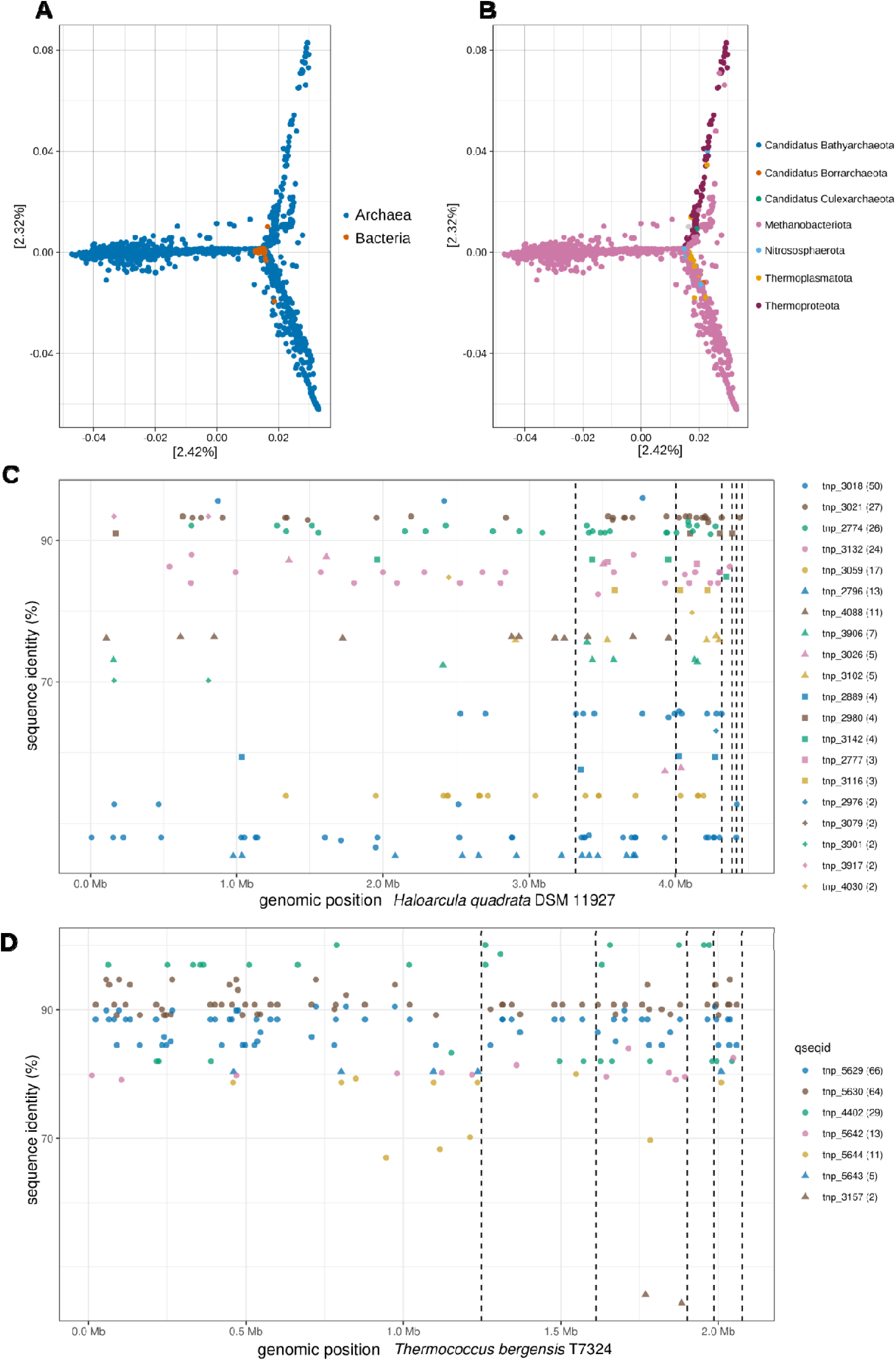
Genomic ordination, taxonomic differentiation, and transposase families across domain-level, phylum-level, and strain-specific contexts. (a) Principal Coordinate Analysis (PCoA) of transposase duplication profiles across archaeal and shared Tnps with bacterial genomes (axis 1: 2.42%, axis 2: 2.32%; PERMANOVA, 999 permutations, R^2^ = 0.00894, p < 0.01). (b) Ordination of archaeal transposase family content at the phylum level, highlighting distinct separation between *Methanobacteriota* and *Thermoproteota*, with intermediate distributions observed for *Thermoplasmatota* and *Nitrosphaerota* (PERMANOVA, 999 permutations, R^2^ = 0.0794, p < 0.01). (c-d) Spatial distribution and sequence identity of duplicated transposases within representative archaeal genomes; individual points represent single hits (n) plotted by cumulative genomic position (X-axis) and amino acid identity (Y-axis), colors and marker shapes denoting distinct Tnp families, with dotted vertical lines distinguishing the main chromosome from extrachromosomal replicons. (c) Genomic distribution and family abundance of Tnps in *Haloarcula quadrata* DSM 11927. (d) Genomic distribution and family abundance of Tnps in *Thermococcus bergensis* T7324. **Alt text. Alt text.** Four-panel figure showing transposase profiles. Panels a and b are PCoA ordination scatter plots comparing transposase content by domain in a and by phylum in b. Panels c and d are scatter plots showing genomic position versus amino acid identity for transposases across chromosomes and plasmids in *Haloarcula quadrata* in panel c and *Thermococcus bergensis* in panel d.

Building on these global patterns, we examined specific duplication events in representative genomes exhibiting high transposase burdens. For example, in *Haloarcula quadrata* DSM 11927, we identified several copies of duplicated Tnps (Figure 4c) distributed across distinct Tnp families throughout the genome in all divergence levels, revealing striking compartmentalization among replicons (Table 3). We observed a higher concentration of these elements within the plasmids, which harbor 120 duplicated hits, than on the main chromosome, despite the main chromosome being significantly larger (∼3.32 Mb).

**Table 3.**
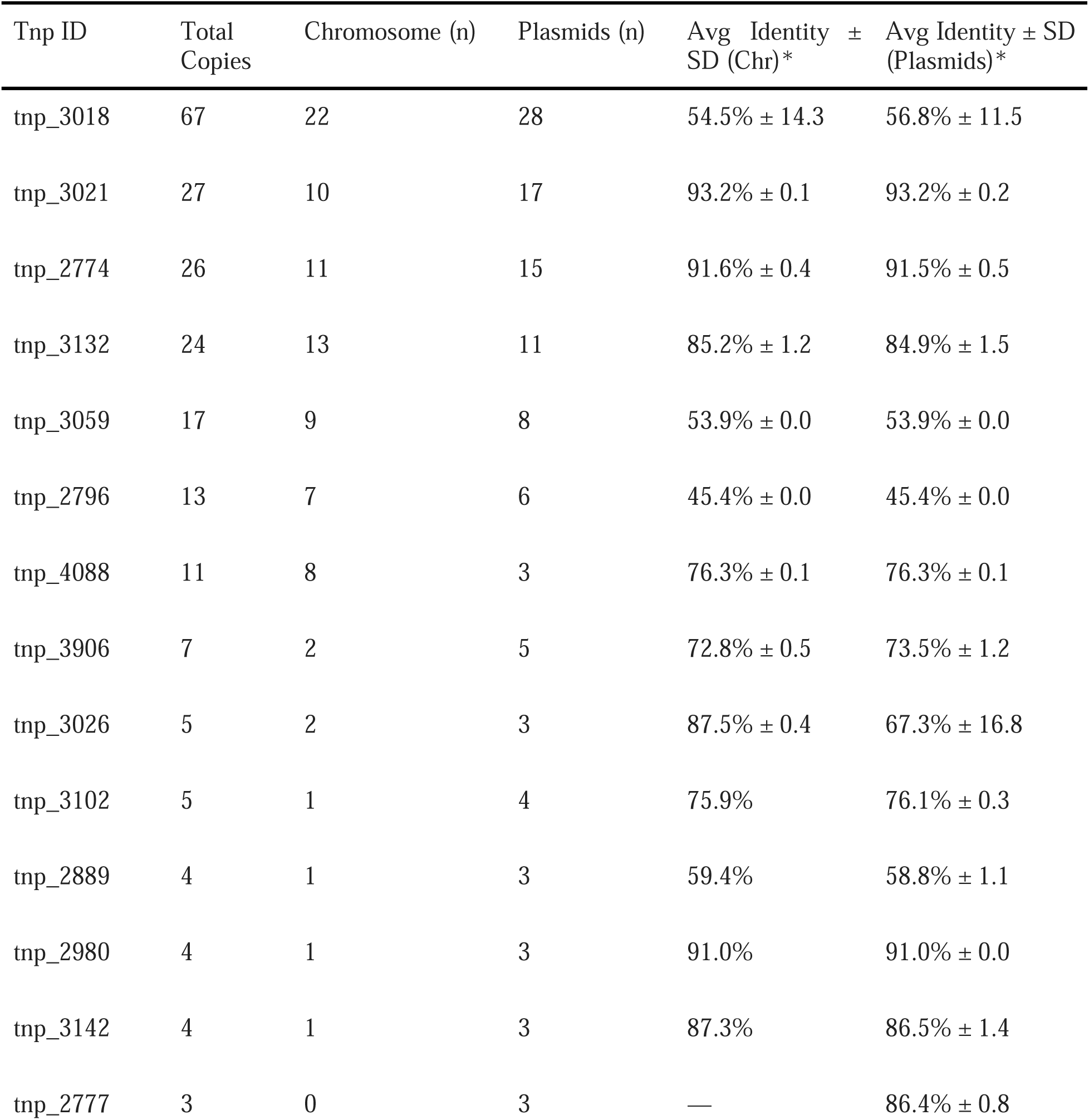
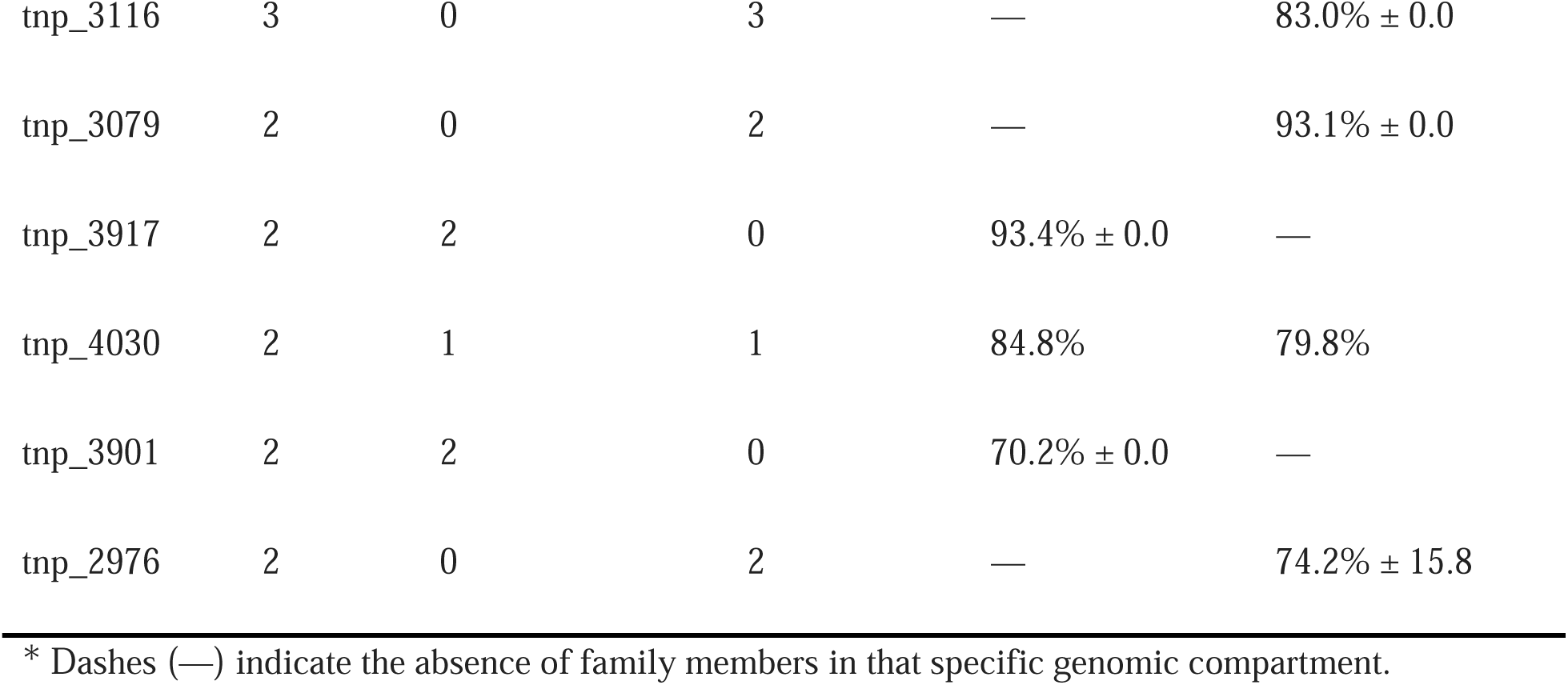
Genomic distribution and sequence identity of duplicated transposase families in *Haloarcula quadrata* DSM 11927. Families are sorted by total abundance (n ≥ 2). Average identity percentages reflect divergence times between the main chromosome and the extrachromosomal fraction (plasmids).

Distinct evolutionary regimes among families strongly shaped this asymmetrical distribution. Highly abundant and conserved families, such as tnp_3021 (27 copies, 93.2% ± 0.2% average identity) and tnp_2774 (26 copies, ∼91.5% average identity; SD ≤ 0.5%), were enriched on plasmids while actively targeting the main chromosome, underscoring contemporary interreplicon transposition dynamics, as evidenced by their remarkably low sequence divergence. Conversely, the single most abundant family, tnp_3018 (67 copies), exhibited a dispersal pattern characterized by low sequence identity and high divergence (54.5% ± 14.3% in chromosomes and 56.8% ± 11.5% in plasmids), indicating a highly heterogeneous, fixed expansion across genomic elements. Furthermore, strict compartmentalization was observed for specific mobile elements, including four families (tnp_2777, tnp_3116, tnp_3079, and tnp_2976) that are confined exclusively to the plasmidial fraction.

Evaluation against evolutionary divergence thresholds (40%, 70%, and 90% identity) indicated that the genomic architecture of *H. quadrata* reflects a multi-tiered history of Tnps accumulation, with recent insertions clustering above 90% identity and older elements in the 40%-70% range.

In contrast, *Thermococcus bergensis* T7324 was analyzed due to its strikingly restricted divergence profile; despite its high total transposase content (Figure 4d), it exhibits a remarkable scarcity of ancient duplications below the 70% sequence identity threshold. While *H. quadrata* sustained heterogeneous families across multiple replicons and evolutionary depths, *T. bergensis* is dominated by a synchronized amplification of the families tnp_4404, tnp_5629, and tnp_5630 (270 of 404 total hits), which show tight identity distributions (∼87% and ∼91%), suggesting that these copies recently multiplied and spread across the genome. Additionally, *H. quadrata* exhibited a more continuous exchange between chromosomal and plasmid-borne transposase populations.

In the domain Bacteria, the transposase distribution was led by class Gammaproteobacteria (181 distinct Tnp families) and family Enterobacteriaceae (134 Tnp families), with maximum intra-species diversity observed in notorious genomes for their metabolic plasticity and open pangenomes, such as *Escherichia coli* (126 families), *Pseudomonas aeruginosa* (103 families), and the nitrogen-fixing soil symbiont *Rhizobium* (68 families). Our screening identified tnp_195 as the most promiscuous Tnp family, spanning 24 distinct bacterial lineages, followed by highly mobile clusters such as tnp_4916, tnp_3056, and tnp_2933 (each present across 23 distinct organisms). The multi-lineage prevalence of these specific clusters suggests they represent highly successful, non-restricted mobilome vectors capable of disseminating across diverse cellular architectures.

To characterize the dynamics of this transfer, we quantified the distribution of the 20 most prevalent shared Tnp families, revealing a distinct gradient of cross-domain promiscuity (Table 4). Our analysis revealed a clear dichotomy in dispersal strategies, with families such as tnp_3691 and tnp_9874 exhibiting extremely high Archaea/Bacteria prevalence ratios (>75), suggesting narrow specialization within the archaeal lineage. In contrast, families such as tnp_3141, tnp_5493, and tnp_1437 show low ratios (<3.5), indicating a much higher capacity to establish and persist across both domains. This distinction suggests that while a significant portion of the archaeal mobilome remains lineage-segregated, a select subset of highly versatile vectors acts as permanent genetic bridges between the disparate genomic architectures of Bacteria and Archaea.

**Table 4.** Distribution profile of the 20 most prevalent shared Transposase (Tnp) families between Archaea and Bacteria. The table shows the prevalence (total organism count) of shared families, highlighting the disparity between Archaea and Bacteria with a calculated Archaea/Bacteria ratio.

| TNP ID | Archaea (n) | Bacteria (n) | Total (n) | Ratio (A/B) |
| --- | --- | --- | --- | --- |
| tnp_6437 | 253 | 9 | 262 | 28.11 |
| tnp_4036 | 158 | 5 | 163 | 31.6 |
| tnp_9874 | 151 | 2 | 153 | 75.5 |
| tnp_9425 | 84 | 15 | 99 | 5.6 |
| tnp_8744 | 95 | 2 | 97 | 47.5 |
| tnp_3691 | 90 | 1 | 91 | 90 |
| tnp_3688 | 88 | 1 | 89 | 88 |
| tnp_3386 | 75 | 3 | 78 | 25 |
| tnp_3387 | 72 | 2 | 74 | 36 |
| tnp_3483 | 72 | 2 | 74 | 36 |
| tnp_3393 | 67 | 4 | 71 | 16.75 |
| tnp_2183 | 56 | 11 | 67 | 5.09 |
| tnp_4594 | 56 | 7 | 63 | 8 |
| tnp_1437 | 44 | 13 | 57 | 3.38 |
| tnp_3527 | 44 | 8 | 52 | 5.5 |
| tnp_4026 | 45 | 3 | 48 | 15 |
| tnp_3571 | 36 | 7 | 43 | 5.14 |
| tnp_3141 | 25 | 11 | 36 | 2.27 |
| tnp_3498 | 35 | 1 | 36 | 35 |
| tnp_5493 | 26 | 9 | 35 | 2.8 |

Extending the comparison to bacteria provides a broader context for understanding the duplication patterns detected in archaeal genomes. Rather than an archaeal-specific phenomenon, MGE expansion appears to be a recurring feature of unusually large, highly duplicated genomes across both domains. A diverse and active mobilome may increase genomic plasticity by enabling DNA rearrangements and the duplication or redistribution of functional blocks, consequently expanding the range of genomic configurations accessible to these organisms. Taken together, these data suggest that the extensive duplication observed in the 48 archaeal outliers may arise, at least in part, from increased mobilome activity and may reflect a broader mechanism of genome expansion shared across Archaea and Bacteria.

## 3. Discussion

Our analysis of 1,829 complete archaeal genomes did not fit with the traditional view that gene duplication is a stochastic, neutral byproduct of genome expansion. Early evolutionary frameworks proposed that gene duplication was the primary mechanism for the emergence of ancient metabolic traits (Jensen 1976; Fani and Fondi 2009); some works have relegated duplicated copies in modern prokaryotic genomes to the status of genomic noise (Krakauer and Nowak 1999; Haubold and Wiehe 2006; Tria and Martin 2021). Contrary to these previous perspectives, our findings showed that paralogous content is a highly regulated, lineage-specific architectural trait. We identify distinct, non-uniform gene duplication patterns across the archaeal domain, evidenced by the lineage-specific accumulation of high-identity duplicates concentrated in specific functional classes-such as mobile elements in *Halobacteria* (Table 3, Figure 4c) and *Thermococci* (Figure 4d), ABC-2 and MFS transporters in Microcaldota and Methanonatronarchaeia (*Persistence index*, PI = 39.12% and PI = 31.22% respectively; Supplementary Tables S2 and S3), and central metabolic enzymes (e.g. like *acetyl-CoA synthetase, K01895*; Supplementary table S4). These metabolic and transport expansions are present in small-genome high-duplication outliers like *Ferroplasma*, *Thermococcus* and *Methanotorris,* which dedicate up to 18.3% of their proteomes to paralogous groups (Supplementary Table S1). These taxa form distinct duplication hotspots with Z-scores > 2 (Figure 2 and Supplementary Figure S1), showing that high-identity gene duplication is lineage-specific rather than a consequence of overall genome growth. Our results indicate that while genome size contributes to the total number of paralogs, it is not a universal predictor of paralogous gene content across the domain Archaea (Figure 1c and Supplementary Table S1). This observation agrees with observations in Bacteria (Pushker et al. 2004), which challenge the assumption that larger genomes necessarily accumulate paralogs linearly with their size (Konstantinidis and Tiedje 2004; Pushker et al. 2004; Grilli et al. 2012; Wang et al. 2025).

Our data highlight the coexistence of compact, highly duplicated genomes with large genomes that exhibit either extensive paralog expansions or surprisingly low duplication levels; this suggests that the accumulation of paralogs is more strongly governed by ecological niche constraints and lifestyle-specific selection pressures than by genome size (Altermann 2012; Dutta and Paul 2012; Salwan and Sharma 2022). Consequently, our findings support a model in which the genomic repertoire of an archaeon is not merely a collection of genes but a balanced investment strategy, in which gene duplication serves as an adaptive trait that contributes to the organism’s stability in its specific environment.

Our identification of specific ’outlier’ genomes, lineages exhibiting disproportionately high levels of paralogous expansion, provides a view into the adaptive limits of Archaea, in which these genomes do not merely accumulate genetic material; they appear to maintain a state of ’permanent duplication’ that stands in stark contrast to the low duplication levels observed in related taxa (Kellner et al. 2018). However, extreme paralogous expansion is only one of multiple viable evolutionary trajectories, and the prevalence of genomes with minimal gene duplications across the archaeal domain underscores that genomic minimization is an equally successful strategy (Csurös and Miklós 2009), likely favored in stable or nutrient-limited environments where maintaining large paralogous repertoires imposes a significant metabolic demand (Baker et al. 2024). Therefore, we propose that the ’outliers’ represent a specialized genomic configuration selected for its ability to protect against extreme selective pressures. Consequently, our analysis suggests that paralog-enriched architectures reflect a high-cost, high-reward strategy for niche-specific colonization. In contrast, non-duplicated architectures reflect a distinct evolutionary investment in metabolic efficiency and optimization.

Our systematic analysis of paralogous content across 28 taxonomic classes reveals a dichotomy, ranging from the ’Permanent Duplication’ of members of Halobacteria to the ’Genomic Stability’ of Nanobdellia, suggesting that archaeal lineages occupy distinct adaptive zones defined by how they manage their paralogous repertoires (Spang et al. 2017). The fact that lineages such as Halobacteria maintain high Z-scores across all identity thresholds indicates that their genomic architecture is based on a constitutive, high-plasticity model (Ludt and Soppa 2019). This suggests that these organisms maintain a diverse pool of paralogs for long-term ecological success, serving as a genomic reservoir (Baker et al. 2024). Our results demonstrate that the archaeal domain is a mosaic of lineages that balance ancient duplications with newly evolved genes, which aid adaptive innovation. However, it’s important to acknowledge that public repositories such as NCBI RefSeq exhibit a historical sampling bias, in which groups like Methanobacteriota dominate complete genome availability, while groups like DPANN and Asgardarchaota remain underrepresented (Supplementary Table S2). While using reference-grade genomes ensures an artifact-free baseline, domain-wide trends may still be influenced by this taxonomic imbalance, expanding future analyses to include both newly isolated reference sequences and metagenome-assembled genomes (MAGs) are needed to validate these results across archaeal diversity.

Our analysis suggests that archaeal adaptation is mediated by a hierarchical investment in specific protein families (Matarredona et al. 2020; Qi et al. 2021; Saini et al. 2023). The observed transition from old generalist machinery (*e.g.*, replication and repair mechanisms) to niche-specific contemporary innovations (*e.g.*, metabolic gene expansions) illustrates a clear decoupling of evolutionary timescales within the proteome (Cummins et al. 2022). The conservation of transcriptional scaffolds such as TFIIB (K03124) and central carbon metabolism enzymes (*e.g.*, acetyl-CoA synthetase) indicates that old duplication events have been repurposed to confer structural robustness to the metabolic core. This layer acts as a genomic foundation, ensuring cellular viability across diverse conditions against metabolic perturbations (Tenorio-Salgado et al. 2024; John et al. 2026).

As the threshold approaches 70%, the duplication repertoire shifts toward reinforcing cellular integrity through high-affinity metal transporters (*e.g.*, *copB, zntA*) and protein-quality-control chaperones (VCP/CDC48), which act as geochemical stressors. This suggests that intermediate paralogs are essential for maintaining physiological homeostasis in environments characterized by fluctuating nutrient availability or chemical gradients (Cisneros et al. 2026).

Recent duplications reveal an expansion of proteins involved in genome protection (*e.g*., SSH7, resolvases) and perhaps niche-specific nutrient acquisition (*e.g*., methylamine-corrinoid methyltransferases). The proliferation of cold-shock protein A (CspA) may indicate that high-identity clusters are driven by immediate adaptive requirements for thermal stability and rapid-response metabolism, and that this strategy may allow organisms to acquire new adaptive traits without altering existing metabolic routes (Mehendiratta et al. 2026).

The persistence of COG unknown functions (category S) across all identity thresholds underscores that our understanding of the archaeal adaptive repertoire remains incomplete. The high sequence identity of these uncharacterized clusters suggests they are active components of the archaeal ’dark’ genome repertoire and represent high-priority targets for future experimental characterization (Pavlopoulos et al. 2023).

Our comprehensive screening of 2,034 transposase (Tnp) families demonstrates that the duplication-enriched ’outlier’ status observed in specific archaeal lineages is a byproduct of a highly active, domain-crossing transposition machinery (Tokuda and Shintani 2024; Peng et al. 2026). The distinct evolutionary trajectories of Tnp families, revealed by our multi-scale identity mapping, uncover a sophisticated strategy of genomic compartmentalization that transcends individual lineages (Rentz et al. 2024; Peng et al. 2026). While the asymmetrical distribution observed in *Haloarcula quadrata*, where high-identity transposase duplicates (*e.g.*, tnp_3021, tnp_2774) are partitioned into extrachromosomal replicons, serves as a clear model for chromosome/plasmid segregation of Tnps, similar dynamics are mirrored in other duplicated-enriched clades. Specifically, in *Thermococcus bergensis* and various members of Methanonatronarchaeia, we observe a comparable tendency to harbor recent transposase duplications within plasmidial fractions. This may suggest that relocating transposases to extrachromosomal replicons is a systemic architectural solution across diverse archaeal outliers (Liu et al. 2025; Ostermayer et al. 2025). By sequestering active transposition machinery in plasmids, these archaea could potentially create a rapid-response reservoir that facilitates niche adaptation without incurring the metabolic load or deleterious fitness costs associated with widespread chromosomal rearrangements (Romaniuk et al. 2018; Valentin-Alvarado et al. 2024). Furthermore, the identification of 153 shared Tnp families between Archaea and Bacteria, with variable Archaea/Bacteria prevalence ratios, suggests a complex model of horizontal gene transfer (HGT) that remains to be explored (Siguier et al. 2015; Cuecas et al. 2017). The presence of these 153 shared Tnp families provides evidence for both historical and ongoing HGT events between the two domains of life, suggesting that these specific MGEs are highly versatile and can cross deep phylogenetic barriers. We identify a series of Tnp families that remain lineage-restricted, but a subset of shared Tnps appears to function as genetic bridges across Domains. This cross-domain mobility underscores that the ’outlier’ genomic architecture is not fully autonomous but is part of a fluid, domain-wide exchange network (Haudiquet et al. 2022; Zhaxybayeva and Nesbø 2026). Our findings suggest that the mobilome is not an accessory component; rather, it serves as a catalyst for the genomic scaling observed in highly duplicated archaeal genomes. These mobile elements may increase the adaptive capacity of the metabolic repertoire by providing the substrates that reorganize metabolic flux and duplicate functional modules (Ma et al. 2023). This dynamic interplay among old, intermediate, and recent thresholds may provide the modularity and robustness required for archaeal lineages to successfully colonize the most fluctuating environments on Earth (Salwan and Sharma 2022; Peters et al. 2026). This aligns with recent large-scale surveys identifying lineage-specific expansions of transposases in extremophilic groups, where the proliferation of specific insertion sequences is frequently coupled with the duplication of environment-related metabolic functions (Peng et al. 2026).

Paralog accumulation across the domain Archaea appears to be a lineage-specific trait shaped by metabolic requirements, transport demands, and mobile element dynamics, rather than a direct consequence of proteome enlargement. While our selection of 1,829 high-quality RefSeq genomes provided a curated, artifact-free baseline for mapping these duplication patterns, expanding future analyses to uncultivated lineages from massive metagenomic datasets will test the universal boundaries of these dynamics. Moreover, the precise functional roles of many uncharacterized duplicated genes remain to be determined, and the high prevalence of these uncharacterized duplications represents a major gap in our knowledge and suggests that Archaea harbor previously unrecognized biochemical pathways that could be exploited for novel applications. Further experimental work investigating how extrachromosomal elements segregate and transfer duplicated genes across domains could reveal fundamental mechanisms of horizontal inheritance. Ultimately, a broader genomic sampling combining long-read pangenomics, structural predictions, and high-throughput functional assays will be necessary to clarify how these duplicated repertoires contribute to archaeal adaptation in diverse and changing environments.

## 4. Materials and Methods

### Archaea genome dataset and sequence curation

We retrieved 1,829 complete archaeal genomes from the NCBI RefSeq database (as of December 2025) via *rsync*, these assemblies correspond to unique NCBI RefSeq complete genome entries (GCF accessions), representing distinct strains and species curated by NCBI. A list of the analyzed genomes is in Supplementary Dataset 2 (see Data Availability Statement). To standardize gene calling and bypass heterogeneous annotation sources, we performed *de novo* protein prediction using Prodigal v2.6.3 (Hyatt et al. 2010) in single-genome mode (Genetic Code 11). Post-prediction, we reformatted all FASTA headers via a custom Perl script (see GitHub project repository) to embed the Genome Assembly Identifier (GCF), ensuring traceability across the pipeline. Taxonomic metadata was standardized using the NCBI Taxonomy database and integrated with genome-level statistics, including total protein count and genome length, generated using EMBOSS *infoseq* (Rice et al. 2000) and custom Bash utilities. The taxonomic curation and hierarchical parsing were performed in R (R Core Team 2021), using the *taxize* (Chamberlain and Szöcs 2013; Chamberlain et al. 2020) and *taxizedb* (Chamberlain et al. 2023) packages to interface with the NCBI backbone. We systematically retrieved the full lineage for each GCF identifier, ensuring that every genome was mapped to its corresponding phylum, class, order, family, and, if possible, genus. For lineages with incomplete nomenclature or those designated as *incertae sedis*, we applied recursive assignment logic that defaulted classification to the immediately higher monophyletic rank to prevent data loss.

### Quantification of gene duplication and clustering

We identified gene family expansions and quantified gene duplication using a dual-scale homology clustering approach that integrates global (inter-genomic) and individual (intra-genomic) evolutionary signals. Global clustering was performed by aggregating all predicted proteomes into a single dataset. This aggregate was processed using MMseqs2 (Steinegger and Söding 2017) across three sequence identity thresholds (40%, 70%, and 90%) with a fixed coverage filter of 70% to ensure structural homology. These thresholds were selected based on the previous work of Pushkert et al. (2004), which defined different evolutionary depths ranging from ancient divergent paralogs to recent, nearly identical gene duplications. To achieve high-resolution intra-genomic analysis, we performed an independent all-versus-all self-clustering for each of the 1,829 archaeal proteomes. This individual-scale approach allowed us to quantify intra-genome paralogy for each lineage without the noise from inter-genomic orthology.

The resulting cluster data were processed using custom Python scripts to derive three primary metrics of genomic complexity. First, we calculated the total duplicate count, defined as the sum of all members within a cluster minus the representative sequence (N–1). Second, we determined singleton frequency, identifying proteins with no detectable homologs within the same genome. Finally, we calculated the duplicated proteome fraction (or paralogous gene content(%)), which represents the proportion of the total proteome that is duplicated. This multi-scale framework enabled us to distinguish between widespread gene families and lineage-specific expansions across the entire Domain Archaea.

### Abundance matrix construction and filtering

To enable a systematic comparison across the 1,829 archaeal lineages, we transformed the raw clustering outputs into high-dimensional count matrices for each identity threshold (40%, 70%, and 90%). In these matrices, rows represented individual protein clusters (families) and columns represented the analyzed genomes, with cell values indicating the number of protein copies (paralogs) belonging to a specific cluster in a given genome. To focus our analysis on gene duplication dynamics and paralogous expansion, we implemented a filtering protocol to prioritize clusters with evidence of gene duplication or inter-genomic conservation. These filtered matrices served as the primary input for downstream statistical modeling, including Z-score normalization (explained in the next section) and the identification of significant gene expansion patterns across the Domain Archaea.

### Statistical modeling and outlier detection

We modeled the relationship between total duplicated proteins and proteome size using linear regression to isolate adaptive expansions from background duplication levels. Standardized residuals were calculated for each identity threshold using the *rstandard()* function in R (R Core Team 2021). This function normalizes the raw residuals by accounting for each genome’s leverage, preventing genomes with extreme sizes from disproportionately biasing the error variance. Because these standardized residuals follow a standard normal distribution with a mean of 0 and a standard deviation of 1, they function directly as Z-scores. Consequently, genomes with standardized residual values exceeding ±2.0 were defined as statistically significant outliers, indicating an atypical number of duplicated proteins relative to their genome size. Relative expansion timing was inferred by tracking fluctuations in Z-scores across the 40%, 70%, and 90% scales. This framework categorizes paralog-enriched lineages into four evolutionary trajectories. Permanent Duplication represents consistent residuals across all scales. Emerging Expansions show Z-scores increasing toward the 90% threshold, reflecting recent duplications with high sequence conservation. Historical Divergences exhibit high residuals at 40% that collapse at higher identity values, indicating that ancient events were followed by diversification. Cooling or diverging lineages are identified by a steady decline in residuals, where sequence divergence outpaces the generation of new duplicates.

### Functional annotation

To provide a high-resolution functional context for the identified gene family expansions, we performed an exhaustive annotation of the representative sequences using eggNOG-mapper (Cantalapiedra et al. 2021) v2.1.9 against the eggNOG v5.0 database. Similarity searches were executed via DIAMOND v2 (Buchfink et al. 2021) in sensitive mode to maximize the recovery of remote homologs. This pipeline yielded a multi-layered functional dataset in which each gene family was mapped to a comprehensive suite of identifiers, including Clusters of Orthologous Groups (COGs), Gene Ontology (GO) terms, and KEGG Orthology (KO) identifiers. This multifaceted annotation was important for systematically organizing the archaeal proteome into coherent functional categories.

To isolate signals from categories such as “Information” from the noise of mobile genetic element (MGE) activity, we implemented a keyword-based classification protocol. Gene families associated with the mobilome were identified using a curated lexicon (*e.g*., “transposase”, “integrase”, “IS[0-9]”, “DDE”, “HTH”, “resolvase”) and categorized separately into “Mobile Elements” for downstream analysis.

Following the initial annotation, we integrated the functional data into an R-based analytical pipeline. We used the KEGGREST package (Tenenbaum 2017) to query the KEGG database (Kanehisa et al. 2025) and map KO identifiers to higher-level biological structures, such as metabolic pathways and functional modules.

### Transposase database construction and distribution analysis

We constructed a specialized transposase reference database to evaluate the impact of the mobilome on prokaryotic gene duplication levels. Initial seed sequences were retrieved from TnCentral (Ross et al. 2021), aggregating curated data from ISFinder (Siguier 2006). The raw dataset (10,657 proteins) underwent sequence clustering via MMseqs2 (Steinegger and Söding 2017) at a 40% identity threshold and 70% query coverage, yielding 2,086 family representatives. Distribution was initially assessed across 1,829 archaeal genomes and subsequently expanded globally to identify domain-specific patterns. This expansion involved an exploratory iTOL-based (Letunic and Bork 2024) evaluation of a bacterial subset to compare transposase family content between domains; we analyzed the relative abundances of these families. Using these relative abundances, we performed a Principal Coordinate Analysis (PCoA) based on Bray-Curtis distances to visualize differences in transposase distribution structure and assessed statistical significance using Permutational Multivariate Analysis of Variance (PERMANOVA) with 999 permutations. Followed by a comprehensive search against the complete NCBI RefSeq database (2025), thereby mitigating sampling bias and capturing rare, lineage-specific mobile elements. We used DIAMOND (Buchfink et al. 2021) v2.0.15 (blastp mode) to execute similarity searches using the 2,086 representative Tnps against a custom-indexed RefSeq database (81,617,152 sequences; 31.2 billion residues). We applied an E-value threshold of 10^-10^ and a minimum query and subject coverage of 70% to ensure functional relevance. Transposase prevalence was calculated as the percentage of unique genera within a lineage harboring specific families. To isolate global dispersal patterns from background noise, we analyzed families detected in >10 unique genera individually and aggregated those detected in fewer genera into a Low Abundance category.

### Visualization and statistical reporting

All statistical analyses and figure generation were conducted in R v4.3.0. Data integration and extensive cleaning of the 1,829-genome dataset were performed using the tidyverse suite, particularly *dplyr* (Wickham et al. 2026), *tidyr* (Wickham et al. 2026), and *stringr* (Wickham 2025), to parse complex taxonomic strings and reshape high-dimensional count matrices. The scaling relationships between proteome size and gene duplication levels were evaluated using Ordinary Least Squares (OLS) linear regression models to identify significant evolutionary deviations. To pinpoint genomes with anomalous duplication levels, we calculated the Studentized residuals for each observation. Genomes exceeding a ±2-standard-deviation threshold from the regression line were categorized as “duplication outliers.” Pearson correlation coefficients (*R*) and coefficients of determination (*R^2^*) were used to quantify the strength of these scaling laws across different taxonomic groups and functional categories. Final visualizations were developed using *ggplot2* (Wickham 2016), leveraging *pheatmap* (Kolde 2025) for hierarchical clustering of transposase distribution and patchwork for assembling multi-panel figures. We prioritized clarity and publication-quality aesthetics, implementing custom color-blind-friendly palettes and refining layouts for high-resolution output. The complete analytical pipeline, including modular R scripts for data tidying, residual calculation, and figure rendering, is publicly available in the project repository https://github.com/genomica-fciencias-unam/Archaea-duplicates.

## 5. Data Availability Statement

Custom computational scripts and pipelines used for data processing and analysis are archived on GitHub https://github.com/genomica-fciencias-unam/Archaea-duplicates. The amino acid sequences of representative members of transposase families matching archaeal homologs, along with associated metadata (Supplementary Dataset 1), have been deposited in Figshare under the Digital Object Identifier (https://doi.org/10.6084/m9.figshare.33301875). All raw proteome datasets analyzed in this work were retrieved from the publicly accessible NCBI RefSeq database (https://www.ncbi.nlm.nih.gov/refseq/), and a list of genome identifiers with their taxonomical information is presented in Supplementary Dataset 2 and deposited in Figshare under the Digital Object Identifier (https://doi.org/10.6084/m9.figshare.33411325).

## Acknowledgments and Funding information

This work was supported by Universidad Nacional Autónoma de México by the projects (DGAPA-PAPIIT-UNAM) IN206824 to L.D.A and (DGAPA-PAPIME-UNAM) PE203625 to I.M.V.. Additional financial support and institutional resources were provided by the Departamento de Biología Celular, Facultad de Ciencias, UNAM. We gratefully acknowledge UNAM ’s “Acuerdos Transformativos Read & Publish” initiative for supporting the Open Access publishing charges for this work. While preparing this work, the authors employed Grammarly and Google Gemini for proofreading and enhancing English phrasing. Following this, they reviewed and edited the manuscript as necessary and assume full responsibility for the final published content.

## Author Contributions

**IM-V:** Conceptualization, Methodology, Formal Analysis, Investigation, Data Curation, Validation, Writing - Original Draft, Writing - Review and Editing, Visualization.

**VH-C:** Formal Analysis, Methodology, Investigation, Data Curation, Writing - Review and Editing.

**GM:** Data Curation, Methodology, Formal Analysis, Validation, Writing - Review and Editing.

**AJ-S:** Data Curation, Methodology, Formal Analysis, Writing - Review and Editing.

**LM-T:** Formal Analysis, Methodology, Validation, Writing - Review and Editing.

**LDA:** Conceptualization, Methodology, Formal Analysis, Investigation, Data Curation, Validation, Writing - Original Draft, Writing - Review and Editing, Visualization, Project administration.

## References

1. Altermann E. Tracing Lifestyle Adaptation in Prokaryotic Genomes. Front Microbiol 2012;3. 10.3389/fmicb.2012.00048.

2. Álvarez-Lugo A, Becerra A. The Role of Gene Duplication in the Divergence of Enzyme Function: A Comparative Approach. Front Genet 2021;12:641817. 10.3389/fgene.2021.641817.

3. Álvarez-Lugo A, Becerra A. The Fate of Duplicated Enzymes in Prokaryotes: The Case of Isomerases. J Mol Evol 2023;91(1):76–92. 10.1007/s00239-022-10085-x.

4. Archibald JM, Roger AJ. Gene duplication and gene conversion shape the evolution of archaeal chaperonins. J Mol Biol 2002;316(5):1041–50. 10.1006/jmbi.2002.5409.

5. Baker BA, Gutiérrez-Preciado A, Rodríguez Del Río Á et al. Expanded phylogeny of extremely halophilic archaea shows multiple independent adaptations to hypersaline environments. Nat Microbiol 2024;9(4):964–75. 10.1038/s41564-024-01647-4.

6. Buchfink B, Reuter K, Drost HG. Sensitive protein alignments at tree-of-life scale using DIAMOND. Nat Methods 2021;18(4):366–8. 10.1038/s41592-021-01101-x.

7. Caetano-Anollés G, Yafremava LS, Gee H et al. The origin and evolution of modern metabolism. Int J Biochem Cell Biol 2009;41(2):285–97. 10.1016/j.biocel.2008.08.022.

8. Cantalapiedra CP, Hernández-Plaza A, Letunic I et al. eggNOG-mapper v2: Functional Annotation, Orthology Assignments, and Domain Prediction at the Metagenomic Scale. Mol Biol Evol 2021;38(12):5825–9. 10.1093/molbev/msab293.

9. Chamberlain S, Szoecs E, Foster Z, Arendsee Z, Boettiger C, Ram K, Bartomeus I, Baumgartner J, O’Donnell J, Oksanen J, Tzovaras BG, Marchand P, Tran V, Salmon M, Li G, Grenié M. Taxize: Taxonomic Information from around the Web., version 0.9.98. R Package. 2020. https://github.com/ropensci/taxize.

10. Chamberlain S, Zebulun Arendsee, Stirling T et al. Ropensci/Taxizedb: Taxizedb v0.3.1, version v0.3.1. Zenodo, 4 Apr. 2023. 10.5281/ZENODO.1158055.

11. Chamberlain SA, Szöcs E. taxize: taxonomic search and retrieval in R. F1000Research 2013;2:191. 10.12688/f1000research.2-191.v2.

12. Cisneros AF, Dibyachintan S, Bédard F et al. Evolutionary causes and consequences of gene duplication. Nat Rev Genet published online 16 Feb. 2026. 10.1038/s41576-026-00935-5.

13. Copley SD. Evolution of new enzymes by gene duplication and divergence. FEBS J 2020;287(7):1262–83. 10.1111/febs.15299.

14. Csurös M, Miklós I. Streamlining and large ancestral genomes in Archaea inferred with a phylogenetic birth-and-death model. Mol Biol Evol 2009;26(9):2087–95. 10.1093/molbev/msp123.

15. Cuecas A, Kanoksilapatham W, Gonzalez JM. Evidence of horizontal gene transfer by transposase gene analyses in Fervidobacterium species. PLOS ONE 2017;12(4):e0173961. 10.1371/journal.pone.0173961.

16. Cummins EA, Hall RJ, McInerney JO et al. Prokaryote pangenomes are dynamic entities. Curr Opin Microbiol 2022;66:73–8. 10.1016/j.mib.2022.01.005.

17. Dutta C, Paul S. Microbial lifestyle and genome signatures. Curr Genomics 2012;13(2):153–62. 10.2174/138920212799860698.

18. Edgell DR, Klenk HP, Doolittle WF. Gene duplications in evolution of archaeal family B DNA polymerases. J Bacteriol 1997;179(8):2632–40. 10.1128/jb.179.8.2632-2640.1997.

19. Edgell DR, Malik SB, Doolittle WF. Evidence of independent gene duplications during the evolution of archaeal and eukaryotic family B DNA polymerases. Mol Biol Evol 1998;15(9):1207–17. 10.1093/oxfordjournals.molbev.a026028.

20. Fani R, Fondi M. Origin and evolution of metabolic pathways. Phys Life Rev 2009;6(1):23–52. 10.1016/j.plrev.2008.12.003.

21. Grilli J, Bassetti B, Maslov S et al. Joint scaling laws in functional and evolutionary categories in prokaryotic genomes. Nucleic Acids Res 2012;40(2):530–40. 10.1093/nar/gkr711.

22. Haubold B, Wiehe T. How repetitive are genomes? BMC Bioinformatics 2006;7(1):541. 10.1186/1471-2105-7-541.

23. Haudiquet M, De Sousa JM, Touchon M et al. Selfish, promiscuous and sometimes useful: how mobile genetic elements drive horizontal gene transfer in microbial populations. Philos Trans R Soc B 2022;377(1861):20210234. 10.1098/rstb.2021.0234.

24. Hooper SD. On the Nature of Gene Innovation: Duplication Patterns in Microbial Genomes. Mol Biol Evol 2003;20(6):945–54. 10.1093/molbev/msg101.

25. Hughes AL. Gene duplication and the origin of novel proteins. Proc Natl Acad Sci 2005;102(25):8791–2. 10.1073/pnas.0503922102.

26. Hyatt D, Chen GL, LoCascio PF et al. Prodigal: prokaryotic gene recognition and translation initiation site identification. BMC Bioinformatics 2010;11(1):119. 10.1186/1471-2105-11-119.

27. Jensen RA. Enzyme recruitment in evolution of new function. Annu Rev Microbiol 1976;30(1):409–25. 10.1146/annurev.mi.30.100176.002205.

28. John J, Ortiz M, Ramond P et al. Functional redundancy and metabolic flexibility of microbial communities in two Mid-Atlantic bays. ISME Commun 2026;6(1):ycag021. 10.1093/ismeco/ycag021.

29. Kanehisa M, Furumichi M, Sato Y et al. KEGG: biological systems database as a model of the real world. Nucleic Acids Res 2025;53(D1):D672–7. 10.1093/nar/gkae909.

30. Kellner S, Spang A, Offre P et al. Genome size evolution in the Archaea. Emerg Top Life Sci 2018;2(4):595–605. 10.1042/ETLS20180021.

31. Kolde R. Pheatmap: Pretty Heatmaps, version 1.0.13. R Package. 2025.

32. Konstantinidis KT, Tiedje JM. Trends between gene content and genome size in prokaryotic species with larger genomes. Proc Natl Acad Sci 2004;101(9):3160–5. 10.1073/pnas.0308653100.

33. Krakauer DC, Nowak MA. Evolutionary preservation of redundant duplicated genes. Semin Cell Dev Biol 1999;10(5):555–9. 10.1006/scdb.1999.0337.

34. Letunic I, Bork P. Interactive Tree of Life (iTOL) v6: recent updates to the phylogenetic tree display and annotation tool. Nucleic Acids Res 2024;52(W1):W78–82. 10.1093/nar/gkae268.

35. Liu H, Sun J, Si J et al. Unexplored diversity and potential functions of extra-chromosomal elements. mSystems 2025;10(9):e00175–25. 10.1128/msystems.00175-25.

36. Ludt K, Soppa J. Polyploidy in halophilic archaea: regulation, evolutionary advantages, and gene conversion. Biochem Soc Trans 2019;47(3):933–44. 10.1042/BST20190256.

37. Ma H, Wang M, Zhang YE et al. The power of “controllers”: Transposon-mediated duplicated genes evolve towards neofunctionalization. J Genet Genomics 2023;50(7):462–72. 10.1016/j.jgg.2023.04.003.

38. Manzano-Morales S, Gabaldon T. Phylogenomics of Asgard archaea reveals a unique blend of prokaryotic-like horizontal transfer and eukaryotic-like gene duplication. Preprint, 25 July 2025. 10.1101/2025.07.24.666526.

39. Martínez-Núñez MA, Poot-Hernandez AC, Rodríguez-Vázquez K et al. Increments and Duplication Events of Enzymes and Transcription Factors Influence Metabolic and Regulatory Diversity in Prokaryotes. PLoS ONE 2013;8(7):e69707. 10.1371/journal.pone.0069707.

40. Matarredona L, Camacho M, Zafrilla B et al. The Role of Stress Proteins in Haloarchaea and Their Adaptive Response to Environmental Shifts. Biomolecules 2020;10(10):1390. 10.3390/biom10101390.

41. Mehendiratta S, Badhwar A, Banerjee A et al. Effect of extreme temperatures on microbial growth and associated activities. In: Decoding Plant–Environment–Microbiome Interactions in Stress-Resilient Agriculture. n.p.: Elsevier, 2026, 51–63. 10.1016/B978-0-443-32932-6.00024-2.

42. Ostermayer J, Guzzi N, Czarnecki J et al. Taming wild replicons: evolution and domestication of large extrachromosomal replicons. Curr Opin Microbiol 2025;88:102657. 10.1016/j.mib.2025.102657.

43. Pavlopoulos GA, Baltoumas FA, Liu S et al. Unraveling the functional dark matter through global metagenomics. Nature 2023;622(7983):594–602. 10.1038/s41586-023-06583-7.

44. Pecoraro V, Zerulla K, Lange C et al. Quantification of Ploidy in Proteobacteria Revealed the Existence of Monoploid, (Mero-)Oligoploid and Polyploid Species. PLoS ONE 2011;6(1):e16392. 10.1371/journal.pone.0016392.

45. Peng XY, Gong ZC, Mao SQ. Comprehensive profiling of insertion sequences and eukaryotic-like transposons across archaeal genomes. BMC Genomics published online 14 May 2026. 10.1186/s12864-026-12948-9.

46. Peters J, LoRicco JG, Saracco M et al. Stability and Adaptation of Proteins, Membranes and Cells to Extreme Temperature and Pressure. J Mol Biol 2026;438(4):169477. 10.1016/j.jmb.2025.169477.

47. Pushker R, Mira A, Rodríguez-Valera F. Comparative genomics of gene-family size in closely related bacteria. Genome Biol 2004;5(4):R27. 10.1186/gb-2004-5-4-r27.

48. Qi YL, Evans PN, Li YX et al. Comparative Genomics Reveals Thermal Adaptation and a High Metabolic Diversity in “ *Candidatus* Bathyarchaeia”. mSystems 2021;6(4):10.1128/msystems.00252-21. 10.1128/msystems.00252-21.

49. R Core Team. R: A Language and Environment for Statistical Computing. Viena, Austria: R Foundation for Statistical Computing, 2021. https://www.r-project.org/.

50. Rentz L, Gehlert FO, Schmitz RA. Genomic Mobility: Transposons in Methanoarchaea. In: Methanogens - Unique Prokaryotes. n.p.: IntechOpen, 2024. 10.5772/intechopen.1007828.

51. Rice P, Longden I, Bleasby A. EMBOSS: The European Molecular Biology Open Software Suite. Trends Genet 2000;16(6):276–7. 10.1016/S0168-9525(00)02024-2.

52. Romaniuk K, Golec P, Dziewit L. Insight Into the Diversity and Possible Role of Plasmids in the Adaptation of Psychrotolerant and Metalotolerant Arthrobacter spp. to Extreme Antarctic Environments. Front Microbiol 2018;9:3144. 10.3389/fmicb.2018.03144.

53. Ross K, Varani AM, Snesrud E et al. TnCentral: a Prokaryotic Transposable Element Database and Web Portal for Transposon Analysis. mBio 2021;12(5):e0206021. 10.1128/mBio.02060-21.

54. Saini A, Kumar A, Singh G et al. Survival Strategies and Stress Adaptations in Halophilic Archaebacteria. In: Dhiman SS, Gnimpieba EZ, Gadhamshetty V (eds), ACS Symposium Series. Washington, DC: American Chemical Society, 2023, 1434.1–21. 10.1021/bk-2023-1434.ch001.

55. Salwan R, Sharma V. Genomics of prokaryotic extremophiles to unfold the mystery of survival in extreme environments. Microbiol Res 2022;264:127156. 10.1016/j.micres.2022.127156.

56. Sheridan PO, Raguideau S, Quince C et al. Gene duplication drives genome expansion in a major lineage of Thaumarchaeota. Nat Commun 2020;11(1):5494. 10.1038/s41467-020-19132-x.

57. Siguier P. ISfinder: the reference centre for bacterial insertion sequences. Nucleic Acids Res 2006;34(90001):D32–6. 10.1093/nar/gkj014.

58. Siguier Patricia, Gourbeyre E, Varani A et al. Everyman’s Guide to Bacterial Insertion Sequences. Microbiol Spectr 2015;3(2):3.2.25. 10.1128/microbiolspec.MDNA3-0030-2014.

59. Soppa J. Polyploidy and community structure. Nat Microbiol 2017;2(2):16261. 10.1038/nmicrobiol.2016.261.

60. Spang A, Caceres EF, Ettema TJG. Genomic exploration of the diversity, ecology, and evolution of the archaeal domain of life. Science 2017;357(6351):eaaf3883. 10.1126/science.aaf3883.

61. Steinegger M, Söding J. MMseqs2 enables sensitive protein sequence searching for the analysis of massive data sets. Nat Biotechnol 2017;35(11):1026–8. 10.1038/nbt.3988.

62. Tenenbaum D. KEGGREST. Bioconductor, 2017. 10.18129/B9.BIOC.KEGGREST.

63. Tenorio-Salgado S, Villalpando-Aguilar JL, Hernandez-Guerrero R et al. Exploring the enzymatic repertoires of Bacteria and Archaea and their associations with metabolic maps. Braz J Microbiol 2024;55(4):3147–57. 10.1007/s42770-024-01462-3.

64. Tokuda M, Shintani M. Microbial evolution through horizontal gene transfer by mobile genetic elements. Microb Biotechnol 2024;17(1):e14408. 10.1111/1751-7915.14408.

65. Tria FDK, Brueckner J, Skejo J et al. Gene Duplications Trace Mitochondria to the Onset of Eukaryote Complexity. Genome Biol Evol 2021;13(5):evab055. 10.1093/gbe/evab055.

66. Tria FDK, Martin WF. Gene Duplications Are At Least 50 Times Less Frequent than Gene Transfers in Prokaryotic Genomes. Genome Biol Evol 2021;13(10):evab224. 10.1093/gbe/evab224.

67. Valentin-Alvarado LE, Shi LD, Appler KE et al. Complete genomes of Asgard archaea reveal diverse integrated and mobile genetic elements. Genome Res 2024;34(10):1595–609. 10.1101/gr.279480.124.

68. Vosseberg J, Van Hooff JJE, Marcet-Houben M et al. Timing the origin of eukaryotic cellular complexity with ancient duplications. Nat Ecol Evol 2020;5(1):92–100. 10.1038/s41559-020-01320-z.

69. Wang P, Guo Q, Jiang X et al. Deciphering gene redundancy in prokaryotic genomes provides evolutionary insights for pathogenicity and its roles in clinical infections. Nat Commun 2025;16(1):10797. 10.1038/s41467-025-65840-7.

70. Wickham H. Ggplot2: Elegant Graphics for Data Analysis. R Package. 2016. https://ggplot2.tidyverse.org.

71. Wickham H. Stringr: Simple, Consistent Wrappers for Common String Operations., version version 1.6.0. R Package. 2025. https://stringr.tidyverse.org.

72. Wickham H, François R, Henry L et al. Dplyr: A Grammar of Data Manipulation., version version 1.2.1. R Package. 2026. https://dplyr.tidyverse.org.

73. Wickham H, Vaughan D, Girlich M. Tidyr: Tidy Messy Data., version version 1.3.2. R Package. 2026. https://tidyr.tidyverse.org.

74. Zhaxybayeva O, Nesbø CL. Impact of Horizontal Gene Transfer on Adaptations to Extreme Environments. J Mol Biol 2026;438(4):169403. 10.1016/j.jmb.2025.169403.

