## Supplementary Material for "Exploring protein duplication across Archaea genomes"

**Supplementary Table S1. Detailed genomic metrics for archaeal lineages identified as gene-duplication outliers.** This table presents the 75 genomes that exceeded the statistical thresholds for gene duplication at 40% sequence identity. Lineages are categorized based on the relationship between their proteome size and duplication metrics: (1) Double Outliers, representing genomes in the top 5th percentile for both absolute duplicate counts and duplicated proteome percentage; and (2) Small-Genome High-Duplication (SGHD), representing genomes with total protein counts below the dataset median (2,880 proteins) that nonetheless fall within the top 5th percentile for duplicated proteome percentage.

| IDs | Taxonomy | Proteome Size | N° of Duplicates at 40% | Duplicated Proteome (%) at 40%* | Category |
| --- | --- | --- | --- | --- | --- |
| GCF_000730175 | *Haloferax sp. ATB1* | 4381 | 1289 | 29.42 | Double Outlier |
| GCF_005435185 | *Halorubrum sp. SD626R* | 6878 | 1866 | 27.13 | Double Outlier |
| GCF_020700235 | *Haladaptatus pallidirubidus YIM 93656* | 6785 | 1773 | 26.13 | Double Outlier |
| GCF_020618475 | *Haladaptatus halobius PSR5* | 8143 | 1998 | 24.54 | Double Outlier |
| GCF_003666005 | *Halorubrum sp. Atlit-9R* | 3893 | 879 | 22.58 | Double Outlier |
| GCF_002135005 | *Halorubrum sp. SD612* | 3612 | 791 | 21.90 | Double Outlier |
| GCF_023077655 | *Halorientalis brevis YC89* | 4718 | 1009 | 21.39 | Double Outlier |
| GCF_021083305 | *Haladaptatus sp. DYF46* | 5931 | 1256 | 21.18 | Double Outlier |
| GCF_031165565 | *Natrinema thermotolerans A29* | 4536 | 958 | 21.12 | Double Outlier |
| GCF_019599505 | *Haloarcula nitratireducens F27* | 5328 | 1095 | 20.55 | Double Outlier |
| GCF_005938085 | *Haloterrigena sp. H1* | 4519 | 910 | 20.14 | Double Outlier |
| GCF_000970145 | *Methanosarcina siciliae C2J* | 4706 | 936 | 19.89 | Double Outlier |
| GCF_004799645 | *Natronorubrum bangense JCM 10635* | 4377 | 863 | 19.72 | Double Outlier |
| GCF_020177375 | *Saliphagus infecundisoli YIM 93745* | 4627 | 907 | 19.60 | Double Outlier |
| GCF_020177165 | *Saliphagus infecundisoli YIM 93707* | 4615 | 902 | 19.54 | Double Outlier |
| GCF_003344565 | *Saliphagus sp. LR7* | 4743 | 919 | 19.38 | Double Outlier |
| GCF_031626495 | *Natrinema sp. 1APR25-10V2* | 5577 | 1063 | 19.06 | Double Outlier |
| GCF_002287235 | *Methanosarcina spelaei MC-15* | 4428 | 839 | 18.95 | Double Outlier |
| GCF_000007345 | *Methanosarcina acetivorans C2A* | 4884 | 922 | 18.88 | Double Outlier |
| GCF_028618695 | *Natrinema soli LMG 29247* | 7105 | 1335 | 18.79 | Double Outlier |
| GCF_024494695 | *Natronoarchaeum rubrum GX48* | 4037 | 753 | 18.65 | Double Outlier |
| GCF_029338335 | *Halomontanus rarus TS33* | 5569 | 1038 | 18.64 | Double Outlier |
| GCF_037335815 | *Salinirubellus litoreus SYNS196* | 5259 | 976 | 18.56 | Double Outlier |
| GCF_001723155 | *Haladaptatus sp. W1* | 5836 | 1069 | 18.32 | Double Outlier |
| GCF_024266705 | *Natrinema gelatinilyticum BND6* | 5461 | 998 | 18.28 | Double Outlier |
| GCF_900110465 | *Halogranum amylolyticum CGMCC 1.10121* | 5334 | 966 | 18.11 | Double Outlier |
| GCF_000025325 | *Haloterrigena turkmenica DSM 5511* | 5224 | 946 | 18.11 | Double Outlier |
| GCF_020614375 | *Haladaptatus salinisoli PSR8* | 5180 | 934 | 18.03 | Double Outlier |
| GCF_004116405 | *Halegenticoccus tardaugens SYSU A00711* | 4948 | 884 | 17.87 | Double Outlier |
| GCF_022870485 | *Halococcus dombrowskii H4* | 4065 | 721 | 17.74 | Double Outlier |
| GCF_024136965 | *Haladaptatus sp. AB618* | 4851 | 858 | 17.69 | Double Outlier |
| GCF_013456555 | *Natrinema zhouii YPL30* | 5431 | 958 | 17.64 | Double Outlier |
| GCF_008122205 | *Natronococcus pandeyae LS1_42* | 5205 | 914 | 17.56 | Double Outlier |
| GCF_024136975 | *Haladaptatus sp. AB643* | 4876 | 851 | 17.45 | Double Outlier |
| GCF_029338575 | *Halomarina halobia PSR21* | 4295 | 749 | 17.44 | Double Outlier |
| GCF_030734285 | *Natronococcus wangiae AD-5* | 5141 | 881 | 17.14 | Double Outlier |
| GCF_029688865 | *Natronococcus sp. A-GB7* | 4935 | 843 | 17.08 | Double Outlier |
| GCF_020405225 | *Natrinema sp. DC36* | 5362 | 914 | 17.05 | Double Outlier |
| GCF_025517495 | *Natronoglomus mannanivorans Arc-xg1-1* | 5264 | 897 | 17.04 | Double Outlier |
| GCF_029338515 | *Haloglomus litoreum DT116* | 4644 | 789 | 16.99 | Double Outlier |
| GCF_024138165 | *Halorussus vallis XZYJT10* | 5048 | 852 | 16.88 | Double Outlier |
| GCF_026248685 | *Haladaptatus caseinilyticus ZJ1* | 5062 | 854 | 16.87 | Double Outlier |
| GCF_002572525 | *Natrinema sp. CBA1119* | 5118 | 863 | 16.86 | Double Outlier |
| GCF_017357405 | *Halomontanus rarus KZCA124* | 5478 | 923 | 16.85 | Double Outlier |
| GCF_028747785 | *Halosimplex aquaticum XZYJT29* | 5045 | 846 | 16.77 | Double Outlier |
| GCF_900156425 | *Haladaptatus litoreus CGMCC 1.7737* | 5014 | 839 | 16.73 | Double Outlier |
| GCF_004217335 | *Natrinema hispanicum DSM 18328* | 4358 | 729 | 16.73 | Double Outlier |
| GCF_000970085 | *Methanosarcina siciliae T4/M* | 4232 | 700 | 16.54 | Double Outlier |
| GCF_036584055 | *Halalkalicoccus salilacus FCH27* | 4463 | 735 | 16.47 | Double Outlier |
| GCF_000336715 | *Halococcus thailandensis JCM 13552* | 4366 | 715 | 16.38 | Double Outlier |
| GCF_002844195 | *Halegenticoccus soli SYSU A9-0* | 4565 | 742 | 16.25 | Double Outlier |
| GCF_024228315 | *Halomarina pelagica BND7* | 4679 | 759 | 16.22 | Double Outlier |
| GCF_000230735 | *Natrinema pellirubrum DSM 15624* | 4353 | 706 | 16.22 | Double Outlier |
| GCF_003841465 | *Natrarchaeobius oligotrophus Archt7* | 4499 | 721 | 16.03 | Double Outlier |
| GCF_005239425 | *Natronorubrum halophilum NEN6* | 4595 | 736 | 16.02 | Double Outlier |
| GCF_023028345 | *Halorientalis marina GDY88* | 4829 | 773 | 16.01 | Double Outlier |
| GCF_024218795 | *Halorussus aquaticus XZYJ18* | 4796 | 767 | 15.99 | Double Outlier |
| GCF_005239435 | *Halorussus salinisoli LT46* | 5034 | 805 | 15.99 | Double Outlier |
| GCF_029502005 | *Natrinema salsiterrestre S1CR25-10* | 4740 | 757 | 15.97 | Double Outlier |
| GCF_000970285 | *Methanosarcina horonobensis HB-1 = JCM 15518 HB-1* | 4415 | 705 | 15.97 | Double Outlier |
| GCF_025947005 | *Halocatena marina QDMS1* | 5162 | 824 | 15.96 | Double Outlier |
| GCF_013342135 | *Haloterrigena gelatinilytica SYSU A121-1* | 4733 | 754 | 15.93 | Double Outlier |
| GCF_023913515 | *Natronococcus zhouii CG52* | 4606 | 731 | 15.87 | Double Outlier |
| GCF_000336935 | *Halococcus salifodinae DSM 8989* | 4323 | 686 | 15.87 | Double Outlier |
| GCF_026013925 | *Halocatena marina RDMS1* | 5260 | 834 | 15.86 | Double Outlier |
| GCF_003977755 | *Haloterrigena salifodinae ZY19* | 4781 | 753 | 15.75 | Double Outlier |
| GCF_003841505 | *Natrarchaeobius chitinivorans Archt4T* | 4453 | 701 | 15.74 | Double Outlier |
| GCF_023008545 | *Natribaculum luteum IBRC-M 10912* | 4624 | 725 | 15.68 | Double Outlier |
| GCF_008728975 | *Haloarcula sp. CBA1131* | 4424 | 693 | 15.66 | Double Outlier |
| GCF_051414035 | *Halosimplex amylolyticum DYHT-AS-1* | 5126 | 802 | 15.65 | Double Outlier |
| GCF_000376445 | *Haladaptatus paucihalophilus DX253* | 4391 | 687 | 15.65 | Double Outlier |
| GCF_023276375 | *Halocatena marina DFN5* | 5438 | 848 | 15.59 | Double Outlier |
| GCF_000496155 | *Ferroplasma sp. Type II na* | 2869 | 541 | 18.86 | SGHD |
| GCF_020386975 | *Thermococcus bergensis T7324* | 2420 | 454 | 18.76 | SGHD |
| GCF_047915675 | *Methanotorris formicicus Mc-S-70* | 2197 | 402 | 18.30 | SGHD |

*Numbers represent percentages.

**Supplementary Table S2. Global profile of gene duplication across archaeal classes.** Comprehensive summary of duplication metrics for all 28 archaeal classes identified in this study. The table includes the number of genomes per class, mean duplication percentages at three identity thresholds (40%, 70%, 90%), and the calculated Persistence Index (PI).

| phylum | class | genomes | 40% | 70% | 90% | Persistence Index^*^ |
| --- | --- | --- | --- | --- | --- | --- |
| C. Borrarchaeota | C. Borrarchaeia | 1 | 11.8 | 2.94 | 0.37 | 3.14 |
| Methanobacteriota | Halobacteria | 781 | 11.26 | 2.44 | 1.06 | 9.41 |
| C. Culexarchaeota | C. Culexarchaeia | 1 | 10.35 | 4.34 | 2.64 | 25.51 |
| Methanobacteriota | Methanomicrobia | 335 | 9.44 | 2.75 | 1.04 | 11.02 |
| Asgardarchaeota | Lokiarchaeia | 1 | 8.35 | 1.29 | 0.43 | 5.15 |
| C. Bathyarchaeota | C. Bathyarchaeia | 5 | 7.61 | 2.05 | 0.75 | 9.86 |
| Thermoproteota | C. Korarchaeia | 3 | 7.23 | 2.27 | 1.38 | 19.09 |
| Methanobacteriota | Methanobacteria | 176 | 6.87 | 1.56 | 0.65 | 9.46 |
| Nitrososphaerota | Conexivisphaeria | 1 | 6.82 | 2.6 | 1.18 | 17.3 |
| Nitrososphaerota | Nitrososphaeria | 50 | 6.77 | 1.76 | 0.55 | 8.12 |
| Methanobacteriota | Archaeoglobi | 16 | 6.5 | 1.43 | 0.68 | 10.46 |
| Promethearchaeota | Promethearchaeia | 1 | 6.49 | 1.06 | 0.46 | 7.09 |
| Thermoproteota | Thermoprotei | 223 | 5.92 | 1.85 | 1.14 | 19.26 |
| Methanobacteriota | Thermococci | 114 | 5.91 | 1.13 | 0.55 | 9.31 |
| Methanobacteriota | Methanonatronarchaeia | 3 | 5.83 | 2.87 | 1.82 | 31.22 |
| Methanobacteriota | Methanococci | 51 | 5.66 | 1.74 | 0.91 | 16.08 |
| Nanobdellati | Microcaldota | 1 | 5.24 | 2.77 | 2.05 | 39.12 |
| Nitrososphaerota | Nitrososphaerota | 9 | 4.6 | 0.78 | 0.13 | 2.83 |
| Thermoplasmatota | Thermoplasmata | 39 | 4.42 | 1.59 | 1.03 | 23.3 |
| Methanobacteriota | C. Hikarchaeia | 1 | 4.2 | 0.34 | 0.09 | 2.14 |
| Methanobacteriota | C. Methanoliparia | 1 | 3.63 | 0.96 | 0.67 | 18.46 |
| Nanobdellota | Nanobdellia | 1 | 3.31 | 0.41 | 0 | 0 |
| Microcaldota | C. Micrarchaeia | 1 | 2.35 | 0.59 | 0.29 | 12.34 |
| Candidatus Nanohalarchaeota | C. Nanohalobia | 1 | 2.27 | 0.08 | 0 | 0 |
| Thermoplasmatota | C. Poseidoniia | 3 | 2.22 | 0.69 | 0.33 | 14.86 |
| Methanobacteriota | Methanopyri | 4 | 2.11 | 0.43 | 0.05 | 2.37 |
| Nanobdellati | C. Nanohalarchaeota | 2 | 1.85 | 0.04 | 0 | 0 |
| Nanobdellota | C. Nanoarchaeia | 2 | 1.06 | 0.23 | 0.04 | 3.77 |

*The Persistence Index (PI) was calculated as the ratio between recent gene duplication levels (90% identity threshold) and ancestral gene duplication levels (40% identity threshold) for each taxonomic class. This metric represents the proportion of high-identity paralogs that have been maintained relative to the total historical pool of duplicated proteins. The index is expressed as a percentage using the formula: PI = (Perc_90_Recent / Perc_40_Ancient) x 100.

**Supplementary Table S3. The top-most frequent functional descriptions of expanded gene families in duplication outliers.** This table summarizes functional enrichment among global paralog-enriched outliers at three sequence-identity thresholds: 40% ("Old"), 70% ("Intermediate"), and 90% ("Recent"). For each threshold, the top 10 functional descriptions are listed alongside their respective COG super-categories. These results demonstrate a clear hierarchical organization of gene duplications, ranging from ancestral energy metabolism and universal stress responses to contemporary mechanisms for cold-shock adaptation and genome integrity.

| Threshold | COG_Category | Description | Ann_Clust_Count^*^ |
| --- | --- | --- | --- |
| 40% | Information | Transcriptional regulator | 10 |
| 40% | Cellular | COG0589 Universal stress protein UspA and related nucleotide-binding proteins | 9 |
| 40% | Information | Transcriptional regulators | 7 |
| 40% | Metabolism | COG1028 Dehydrogenases with different specificities (related to short-chain alcohol dehydrogenases) | 7 |
| 40% | Information | COG1522 Transcriptional regulators | 6 |
| 40% | Information | Stabilizes TBP binding to an archaeal box-A promoter. Also responsible for recruiting RNA polymerase II to the pre- initiation complex (DNA-TBP-TFIIB) | 6 |
| 40% | Metabolism | ABC-type multidrug transport system, ATPase component | 6 |
| 40% | Metabolism | COG0477 Permeases of the major facilitator superfamily | 6 |
| 40% | Metabolism | COG1960 Acyl-CoA dehydrogenases | 6 |
| 40% | Information | Involved in the regulation of DNA replication | 5 |
| 40% | Metabolism | COG0318 Acyl-CoA synthetases (AMP-forming) AMP-acid ligases II | 5 |
| 70% | Information | Transcriptional regulator | 12 |
| 70% | Information | Stabilizes TBP binding to an archaeal box-A promoter. Also responsible for recruiting RNA polymerase II to the pre- initiation complex (DNA-TBP-TFIIB) | 6 |
| 70% | Information | A general factor that plays a role in the activation of archaeal genes transcribed by RNA polymerase. Binds specifically to the TATA box promoter element, which lies close to the position of transcription initiation | 5 |
| 70% | Information | Transcriptional regulators | 5 |
| 70% | Metabolism | COG2217 Cation transport ATPase | 5 |
| 70% | Poorly Char. | Peptidase_C39-like family | 5 |
| 70% | Metabolism | COG1028 Dehydrogenases with different specificities (related to short-chain alcohol dehydrogenases) | 4 |
| 70% | Cellular | COG0330 Membrane protease subunits, stomatin prohibitin homologs | 3 |
| 70% | Cellular | COG1192 ATPases involved in chromosome partitioning | 3 |
| 70% | Cellular | Transcriptional regulator | 3 |
| 70% | Information | Cold shock | 3 |
| 70% | Information | Seems to be required for the maximal rate of protein biosynthesis. Enhances ribosome dissociation into subunits and stabilizes the binding of the initiator Met-tRNA(I) to 40 S ribosomal subunits | 3 |
| 70% | Metabolism | Belongs to the Glu Leu Phe Val dehydrogenases family | 3 |
| 70% | Metabolism | Belongs to the heme-copper respiratory oxidase family | 3 |
| 70% | Metabolism | Belongs to the sodium solute symporter (SSF) (TC 2.A.21) family | 3 |
| 70% | Metabolism | COG0365 Acyl-coenzyme A synthetases AMP-(fatty) acid ligases | 3 |
| 70% | Metabolism | Major facilitator superfamily | 3 |
| 70% | Poorly Char. | Peptidase domain protein | 3 |
| 70% | Poorly Char. | TRAM domain | 3 |
| 70% | Poorly Char. | zinc finger | 3 |
| 90% | Information | Transcriptional regulator | 8 |
| 90% | Cellular | Peptidase A24A, prepilin type IV | 6 |
| 90% | Metabolism | COG0318 Acyl-CoA synthetases (AMP-forming) AMP-acid ligases II | 6 |
| 90% | Cellular | COG0719 ABC-type transport system involved in Fe-S cluster assembly, permease component | 5 |
| 90% | Information | COG1278 Cold shock proteins | 5 |
| 90% | Metabolism | methyltransferase activity | 5 |
| 90% | Information | TIGRFAM DNA binding domain, excisionase family | 4 |
| 90% | Poorly Char. | Peptidase_C39-like family | 4 |
| 90% | Cellular | COG1192 ATPases involved in chromosome partitioning | 3 |
| 90% | Cellular | Motif C-terminal to PAS motifs (likely to contribute to PAS structural domain) | 3 |
| 90% | Cellular | Transcriptional regulator | 3 |
| 90% | Information | COG1599 Single-stranded DNA-binding replication protein A (RPA), large (70 kD) subunit and related ssDNA-binding proteins | 3 |
| 90% | Information | Can constrain negative DNA supercoils. May be involved in maintaining the integrity of the genome at high temperatures | 3 |
| 90% | Information | Cold shock | 3 |
| 90% | Information | Domain of unknown function (DUF4277) | 3 |
| 90% | Metabolism | Dimethylamine methyltransferase (Dimeth_PyL) | 3 |
| 90% | Metabolism | Major facilitator Superfamily | 3 |
| 90% | Metabolism | PFAM Thiolase | 3 |
| 90% | Metabolism | Part of a complex that catalyzes the reversible cleavage of acetyl-CoA, allowing | 3 |
| 90% | Poorly Char. | ATPase, AAA superfamily | 3 |
| 90% | Poorly Char. | cobalamin binding | 3 |

*The Ann_Clust_Count (Annotated_Cluster_Count) indicates the number of independent protein clusters identified by MMseqs2 that were successfully annotated via eggNOG-mapper. Each cluster contains a variable number of duplicates.

**Supplementary Table S4. Stratified Functional Distribution of KEGG Orthologs Across Evolutionary Scales.** This table summarizes the top 15 most frequent KEGG Orthology (KO) definitions identified in paralog-enriched lineages, categorized by sequence-identity thresholds. Data were filtered to exclude mobile genetic elements (transposases, integrases, and resolvases associated with IS elements) to focus on the metabolic and structural robustness of the outlier taxa.

| Threshold | KEGG ko | KEGG Definition | Occurrence Count |
| --- | --- | --- | --- |
| 40% | ko:K01990 | ABC-2.A; ABC-2 type transport system ATP-binding protein | 7 |
| 40% | ko:K01895 | ACSS1_2, acs; acetyl-CoA synthetase [EC:6.2.1.1] | 6 |
| 40% | ko:K03124 | TFIIB, GTF2B, SUA7, tfb; transcription initiation factor TFIIB | 6 |
| 40% | ko:K00249 | ACADM, acd; acyl-CoA dehydrogenase [EC:1.3.8.7] | 4 |
| 40% | ko:K00666 | K00666; fatty-acyl-CoA synthase [EC:6.2.1.-] | 4 |
| 40% | ko:K02035 | ABC.PE.S; peptide/nickel transport system substrate-binding protein | 4 |
| 40% | ko:K03499 | trkA, ktrA, ktrC; trk/ktr system potassium uptake protein | 4 |
| 40% | ko:K10725 | cdc6A; archaeal cell division control protein 6 | 4 |
| 40% | ko:K00001 | E1.1.1.1, adh; alcohol dehydrogenase [EC:1.1.1.1] | 3 |
| 40% | ko:K00059 | fabG, OAR1; 3-oxoacyl-[acyl-carrier protein] reductase [EC:1.1.1.100] | 3 |
| 40% | ko:K00384 | trxB, TRR; thioredoxin reductase (NADPH) [EC:1.8.1.9] | 3 |
| 40% | ko:K00973 | rfbA, rmlA, rffH; glucose-1-phosphate thymidylyltransferase [EC:2.7.7.24] | 3 |
| 40% | ko:K01784 | galE, GALE; UDP-glucose 4-epimerase [EC:5.1.3.2] | 3 |
| 40% | ko:K01992 | ABC-2.P; ABC-2 type transport system permease protein | 3 |
| 40% | ko:K03406 | mcp; methyl-accepting chemotaxis protein | 3 |
| 70% | ko:K01895 | ACSS1_2, acs; acetyl-CoA synthetase [EC:6.2.1.1] | 6 |
| 70% | ko:K03124 | TFIIB, GTF2B, SUA7, tfb; transcription initiation factor TFIIB | 6 |
| 70% | ko:K03120 | TBP, tbp; transcription initiation factor TFIID TATA-box-binding protein | 5 |
| 70% | ko:K13525 | VCP, CDC48; transitional endoplasmic reticulum ATPase | 4 |
| 70% | ko:K00175 | korB, oorB, oforB; 2-oxoglutarate/2-oxoacid ferredoxin oxidoreductase subunit beta [EC:1.2.7.3 1.2.7.11] | 3 |
| 70% | ko:K00201 | fwdB, fmdB; formylmethanofuran dehydrogenase subunit B [EC:1.2.7.12] | 3 |
| 70% | ko:K00973 | rfbA, rmlA, rffH; glucose-1-phosphate thymidylyltransferase [EC:2.7.7.24] | 3 |
| 70% | ko:K01533 | copB; P-type Cu2+ transporter [EC:7.2.2.9] | 3 |
| 70% | ko:K01534 | zntA; Zn2+/Cd2+-exporting ATPase [EC:7.2.2.12 7.2.2.21] | 3 |
| 70% | ko:K01784 | galE, GALE; UDP-glucose 4-epimerase [EC:5.1.3.2] | 3 |
| 70% | ko:K01912 | paaK; phenylacetate-CoA ligase [EC:6.2.1.30] | 3 |
| 70% | ko:K03236 | EIF1A; translation initiation factor 1A | 3 |
| 70% | ko:K03609 | minD; septum site-determining protein MinD | 3 |
| 70% | ko:K03704 | cspA; cold shock protein | 3 |
| 70% | ko:K07133 | K07133; uncharacterized protein | 3 |
| 90% | ko:K03704 | cspA; cold shock protein | 8 |
| 90% | ko:K16176 | mtmB; methylamine---corrinoid protein Co-methyltransferase [EC:2.1.1.248] | 6 |
| 90% | ko:K07450 | K07450; putative resolvase | 5 |
| 90% | ko:K09014 | sufB; Fe-S cluster assembly protein SufB | 5 |
| 90% | ko:K07133 | K07133; uncharacterized protein | 4 |
| 90% | ko:K16178 | mtbB; dimethylamine---corrinoid protein Co-methyltransferase [EC:2.1.1.249] | 4 |
| 90% | ko:K00666 | K00666; fatty-acyl-CoA synthase [EC:6.2.1.-] | 3 |
| 90% | ko:K16177 | mtmC; monomethylamine corrinoid protein | 3 |
| 90% | ko:K16640 | ssh7; DNA-binding protein 7 [EC:3.1.27.-] | 3 |
| 90% | ko:K00197 | cdhE, acsC; acetyl-CoA decarbonylase/synthase, CODH/ACS complex subunit gamma [EC:2.1.1.245] | 2 |
| 90% | ko:K00201 | fwdB, fmdB; formylmethanofuran dehydrogenase subunit B [EC:1.2.7.12] | 2 |
| 90% | ko:K00626 | ACAT, atoB; acetyl-CoA C-acetyltransferase [EC:2.3.1.9] | 2 |
| 90% | ko:K00721 | DPM1; dolichol-phosphate mannosyltransferase [EC:2.4.1.83] | 2 |
| 90% | ko:K00960 | rpo13; DNA-directed RNA polymerase subunit 13 [EC:2.7.7.6] | 2 |
| 90% | ko:K01895 | ACSS1_2, acs; acetyl-CoA synthetase [EC:6.2.1.1] | 2 |

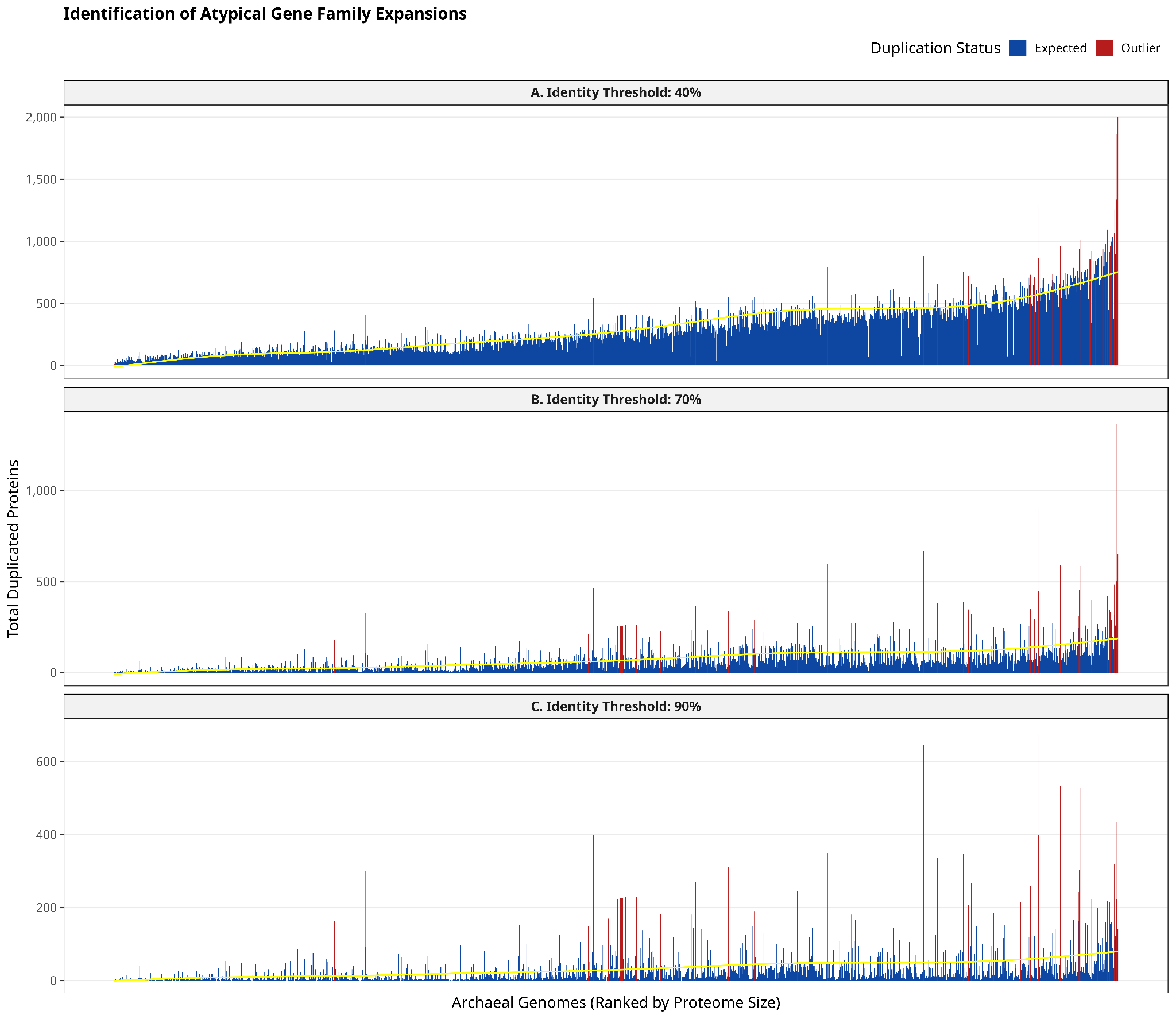

**Supplementary Fig. S1. Statistical identification of atypical gene family expansions across identity thresholds.** (A-C) Distribution of observed versus expected duplicated proteins across 40%, 70%, and 90% sequence identity thresholds. Genomes are ranked along the x-axis by total proteome size. Blue bars represent the observed count of duplicated proteins for each genome, while the yellow line indicates the expected duplication count predicted by the linear model (Duplicates ~ Proteome Size). Genomes with standardized residuals greater than 2 standard deviations (>2 SD) from the expected value are highlighted in red, indicating non-random gene expansion that exceeds the genome-scale baseline.

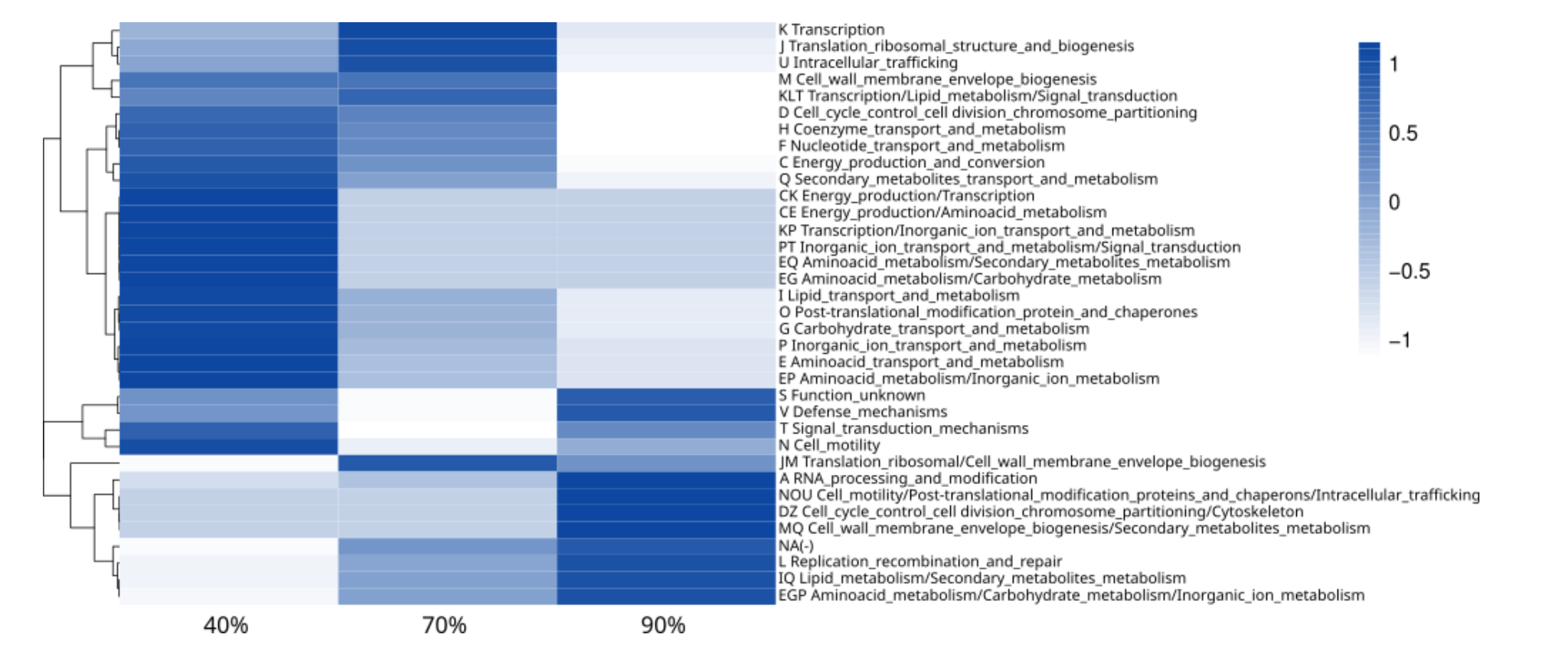

**Supplementary Fig. S2. COG functional distribution of archaeal gene duplications across three identity thresholds.** The heatmap shows the normalized abundance profiles of functional categories across gene duplication events grouped by 40%, 70%, and 90% sequence identity. The color gradient reflects a Z-score transformation, distinguishing enriched biological processes from depleted ones. Functional annotations include single COG codes and complex multi-domain intersections. This comparative view reveals which functional systems exhibited distinct expansion rates across the evolutionary trajectories of the analyzed genomes.

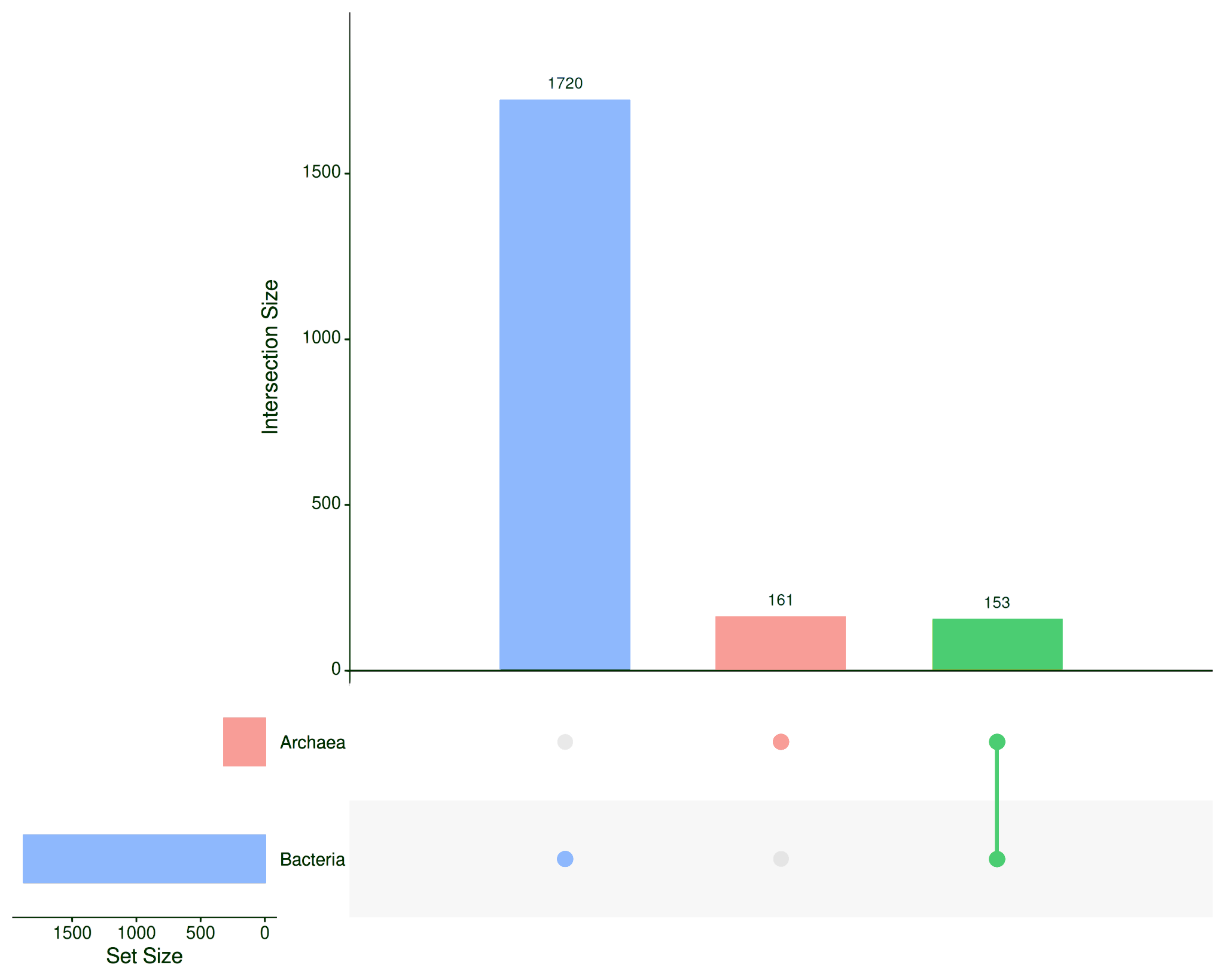

**Supplementary Fig. S3. Distribution of Transposase (Tnp) families across bacterial and archaeal genomes.** A total of 1,720 families are exclusively restricted to Bacteria (green bar). Conversely, 161 families are exclusive to Archaea (blue bar), and 153 families are shared between the two domains (yellow bar).

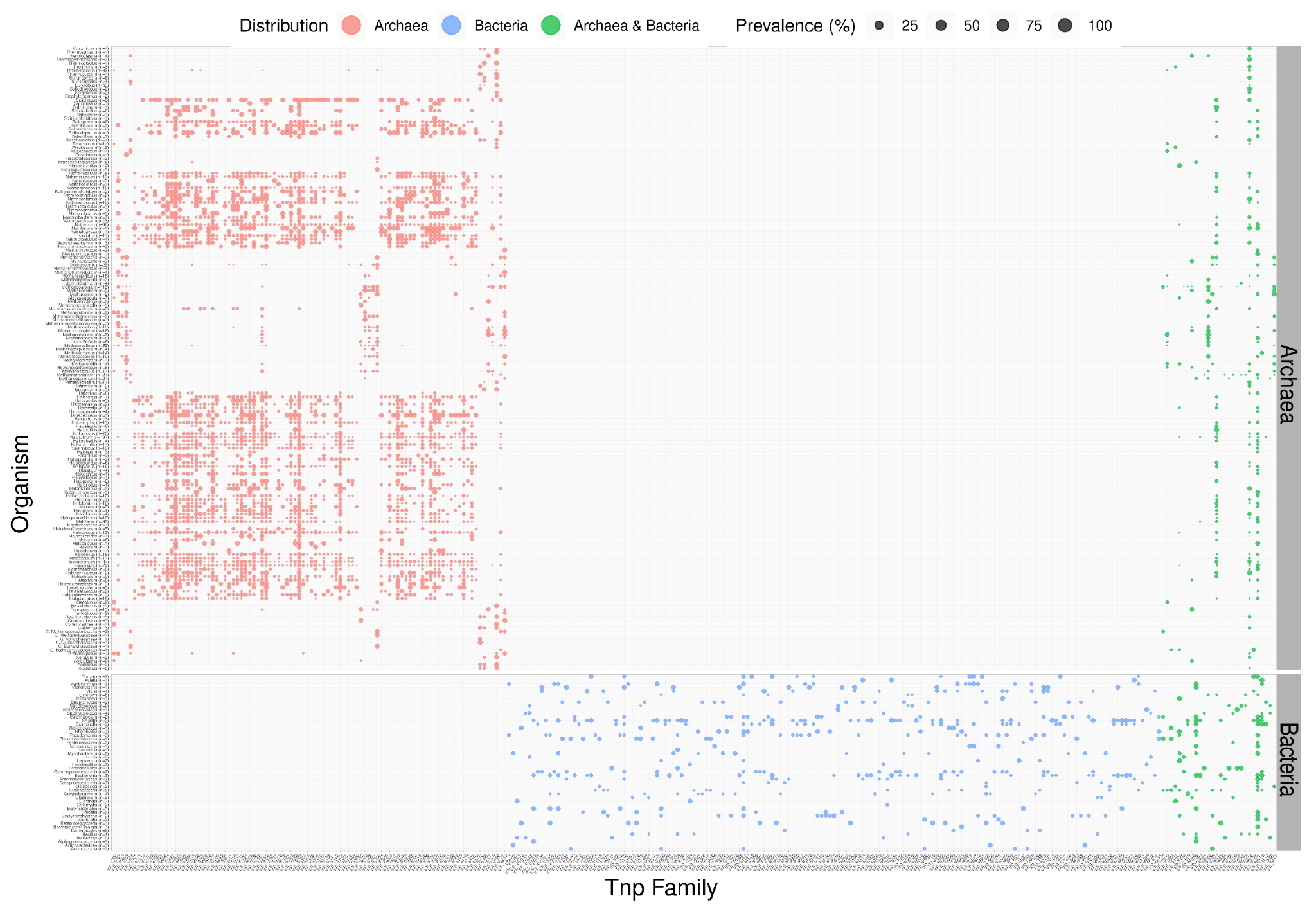

**Supplementary Fig. S4. Comparative prevalence of Transposase (Tnp) families across Archaea and Bacteria.** The dot plot shows the prevalence (the percentage of genomes in a group that contain at least one copy) of representative Tnp families. Genomes are clustered by Domain (Archaea and Bacteria). Dot size is proportional to the prevalence of each Tnp family, ranging from smallest to largest. Only Tnp families present in at least 10 archaeal genera or 2 bacterial genera are shown to highlight widely distributed elements.

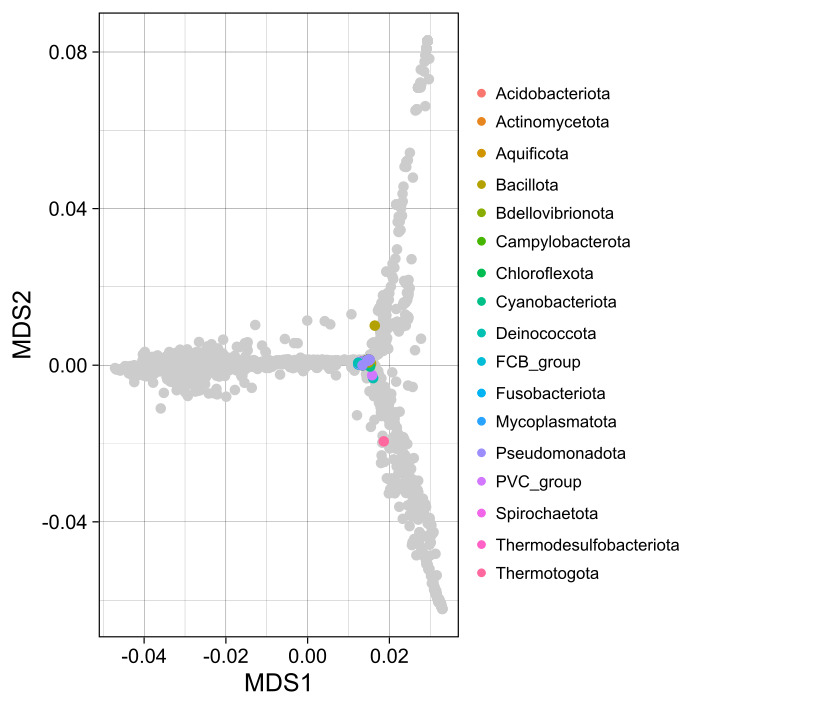

**Supplementary Fig. S5. Principal Coordinate Analysis (PCoA) of transposase (Tnp) family duplication profiles across bacterial phyla.** PCoA based on Bray-Curtis distances illustrating the structural distribution and variation of transposase family content among bacterial genomes. Colored points denote bacterial phyla; corresponding taxonomic groups are specified in the right-hand panel legend.
